# A Layered Microcircuit Model of Somatosensory Cortex with Three Interneuron Types and Cell-Type-Specific Short-Term Plasticity

**DOI:** 10.1101/2023.10.26.563698

**Authors:** Han-Jia Jiang, Guanxiao Qi, Renato Duarte, Dirk Feldmeyer, Sacha J van Albada

## Abstract

Three major types of GABAergic interneurons, parvalbumin- (PV), somatostatin- (SOM) and vasoactive intestinal peptide-expressing (VIP) cells, play critical but distinct roles in the cortical microcircuitry. Their specific electrophysiology and connectivity shape their inhibitory functions. To study the network dynamics and signal processing specific to these cell types in the cerebral cortex, we developed a multi-layer model incorporating biologically realistic interneuron parameters from rodent somatosensory cortex. The model is fitted to *in vivo* data on cell-type-specific population firing rates. With a protocol of cell-type-specific stimulation, network responses when activating different neuron types are examined. The model reproduces the experimentally observed inhibitory effects of PV and SOM cells and disinhibitory effect of VIP cells on excitatory cells. We further create a version of the model incorporating cell-type-specific short-term synaptic plasticity (STP). While the ongoing activity with and without STP is similar, STP modulates the responses of Exc, SOM, and VIP cells to cell-type-specific stimulation, presumably by changing the dominant inhibitory pathways. With slight adjustments, the model also reproduces sensory responses of specific interneuron types recorded *in vivo*. Our model provides predictions on network dynamics involving cell-type-specific short-term plasticity and can serve to explore the computational roles of inhibitory interneurons in sensory functions.

## Introduction

Cortical GABAergic interneurons are inhibitory neurons that modulate and limit the degree of neuronal excitability in the neocortex. They can be classified according to electrophysiological or morphological characteristics or with molecular markers (Ascoli et al., 2008). Although challenges still exist in establishing consistency among different classification methods, many studies in recent years have used molecular markers, which label groups with different genetic origins, to make significant progress in exploring interneuron circuits (Tremblay et al., 2016; Campagnola et al., 2022). The three most common types of cortical interneurons express parvalbumin (PV), somatostatin (SOM), and vasoactive intestinal peptide (VIP), respectively, and play critical but distinct roles in cortical microcircuitry (Karnani et al., 2014; Tremblay et al., 2016; Campagnola et al., 2022). These three types have their own specific electrophysiology, morphology, connectivity, and short-term synaptic plasticity (STP). How these properties are related to their distinct dynamics and inhibitory functions is an important topic for understanding cortical microcircuitry.

PV and SOM cells have been extensively studied and compared experimentally. PV cells are considered a major stabilizing force that produces fast and reliable inhibition, mediated by synaptic activities that are weakened in amplitude (depressed) during high-frequency stimulation (Beierlein et al., 2003; Silberberg and Markram, 2007; Hu et al., 2014; Karnani et al., 2014; Naka and Adesnik, 2016). In contrast, SOM cells act more slowly, and have synaptic effects that are initially weak but can increase in amplitude (facilitate) in response to sustained high-frequency stimulation (Beierlein et al., 2003; Kapfer et al., 2007; Silberberg and Markram, 2007; Karnani et al., 2014; Yavorska and Wehr, 2016). Due to their different synaptic dynamics, PV and SOM cells contribute to cortical sensory processing (Natan et al., 2015, 2017; Seay et al., 2020) and control oscillatory activity (Chen et al., 2017; Van Derveer et al., 2021) in different but complementary ways. On the other hand, VIP cell activity results in disinhibition of pyramidal cells by inhibiting SOM cells (Lee et al., 2013; Pi et al., 2013; Karnani et al., 2016a). Distal or neuromodulatory inputs have been found to alter the activity of VIP cells to mediate disinhibition of sensory signals during different behavioral states, such as wakefulness and movements (Lee et al., 2013; Pi et al., 2013; Fu et al., 2014; Karnani et al., 2016a). Inhibitory interneurons thus form an integral part of the cortical computational circuitry, with different types contributing to different aspects of inhibitory control and complementing each other.

In addition, interneurons show a high degree of diversity across cortical layers. PV, SOM, and VIP cells have different morphologies and connectivities across layers (Xu et al., 2013; Prönneke et al., 2015; Tremblay et al., 2016; Feldmeyer et al., 2018). Furthermore, SOM cells exhibit different target preferences (Xu et al., 2013) and whisking-related activity (Muñoz et al., 2017) across layers of barrel cortex, resulting in layer-specific inhibitory modulation of network activity (Xu et al., 2013; Muñoz et al., 2017). The importance of this cross-layer diversity for computations in the cortical column remains largely unexplored (Hahn et al., 2022).

However, experiments are limited by the number of neurons that can be recorded in each animal or sample. In recent years, there has been a rapid development of genetic methods that allow different types of interneurons to be labeled and studied in living brains or brain slices (Huang and Zeng, 2013; Huang, 2014). But even with genetic labeling, it is still relatively difficult to identify interneurons of a specific type *in vivo*, especially when it is located in deep cortical layers. Furthermore, *in vivo* paired recordings with specific interneuron types are even more technically challenging, which hinders the study of interneuron-type-specific synaptic dynamics. Therefore, to systematically study the roles of interneuron types, computational studies that take the interneuron diversity into account are needed in addition to experimental approaches. Computational models can help to develop a mechanistic understanding of interneuron functions and lead to hypotheses for experimental studies. In particular, understanding interneuron function and dynamics across layers is essential for the study of the multi-layered cortical microcircuitry.

A number of modeling studies have considered the roles of inhibitory neuron types in local cortical circuit dynamics. Yang et al. (2016) and Hertäg and Sprekeler (2019) examined how mutual inhibition between SOM and VIP cells allows switching between two processing modes in which top-down inputs to pyramidal cells are either integrated or canceled. Litwin-Kumar et al. (2016) studied PV, SOM, and VIP cells in firing rate models and revealed their dissociation in firing rate changes in the paradoxical effect, where stimulated inhibitory neurons show a paradoxical decrease in firing rate (Tsodyks et al., 1997; Murphy and Miller, 2009; Ozeki et al., 2009). Mahrach et al. (2020) compared experimental data of PV cell stimulation with their models and found that, compared to a simpler two-population (excitatory and inhibitory) model, a model with the three interneuron types can reflect more details in the experimental data, showing the paradoxical effect whether the network is inhibition-stabilized or not. Bos et al. (2020) further investigated the mechanics of a network with PV and SOM cells and showed how to predict their mean-field behaviors. Lee et al. (2017) and Lee and Mihalas (2017) studied the role of the three interneuron types in single- and multi-column sensory signal processing involving surround suppression and contexual modulation. While these studies have provided theoretical explanations for the functions of PV, SOM and VIP cells, a model that accounts for their diversity across layers is still needed to further understand their roles in a multi-layered cortical column. A few multi-layer models with interneuron types have been developed based on experimental data (Markram et al., 2015; Billeh et al., 2020; Borges et al., 2022; Moreni et al., 2023). However, incorporating the layer- and cell-type-specific electrophysiological properties and synaptic dynamics of an interneuron type into the model while maintaining model simplicity and low simulation cost still remains a challenge, that, if achieved, will allow for more convenient use, adaptation, systematic analysis and generalization of the model.

We therefore developed a new cortical microcircuit model adapted from Potjans and Diesmann (2014), that incorporates PV, SOM and VIP cells. This new model includes layer-specific electrophysiological and synaptic properties of excitatory neurons (Exc) and the three interneuron types. We focus on a mouse barrel column, taking parameters from mouse somatosensory cortex (S1) when available, complementing these with rat S1 data. Like Potjans and Diesmann (2014), we use a common and computationally low-cost neuron model, the Leaky Integrate-and-Fire (LIF) point neuron model. With the LIF neuron model, we can already capture essential aspects of experimentally observed resting-state activity, including cell-type-specific firing rates. To study the effects of synaptic dynamics, we also incorporate cell-type-specific STP parameters derived from experimental data, allowing comparisons between model versions with and without STP. We further use a protocol of cell-type-specific stimulation in L2/3 and L4 to study network responses when different interneuron types are activated. With this protocol, we reproduce experimentally observed inhibitory or disinhibitory effects of PV, SOM, and VIP cells. We further show that STP may modulate population responses to cell-type-specific stimulation, by altering the dominant inhibitory pathways. Specifically, STP qualitatively modifies these responses: (1) Exc responses to Exc stimulation in L2/3; (2) VIP responses to PV stimulation in L2/3; (3) SOM responses to Exc stimulation in L4. We hypothesize and show supporting data that this is because STP affects the following pathways: (1) Exc→SOM→Exc in L2/3; (2) PV→SOM→VIP in L2/3; (3) Exc→PV→SOM in L4. In summary, we created a simple, biologically plausible, and computationally efficient model for the analysis of the roles of interneuron types in the cortical microcircuitry.

## Methods

### Model Overview

Our model is adapted from the multi-layer cortical microcircuit model by Potjans and Diesmann (2014). We extend the model to include three major interneuron types: PV, SOM, and VIP, and use only experimental data on mouse and rat somatosensory cortex. Figure 1 shows an overview of the populations and their synaptic connectivity. All neurons are modeled as Leaky-Integrate-and-Fire (LIF) neurons with exponentially decaying postsynaptic currents (PSCs)^1^ (Tsodyks et al., 2000). Figure 1 shows the dimensions of the model, which correspond to those of a mouse C2 barrel column described previously (Lefort et al., 2009; Petersen, 2019).

**Figure 1.**
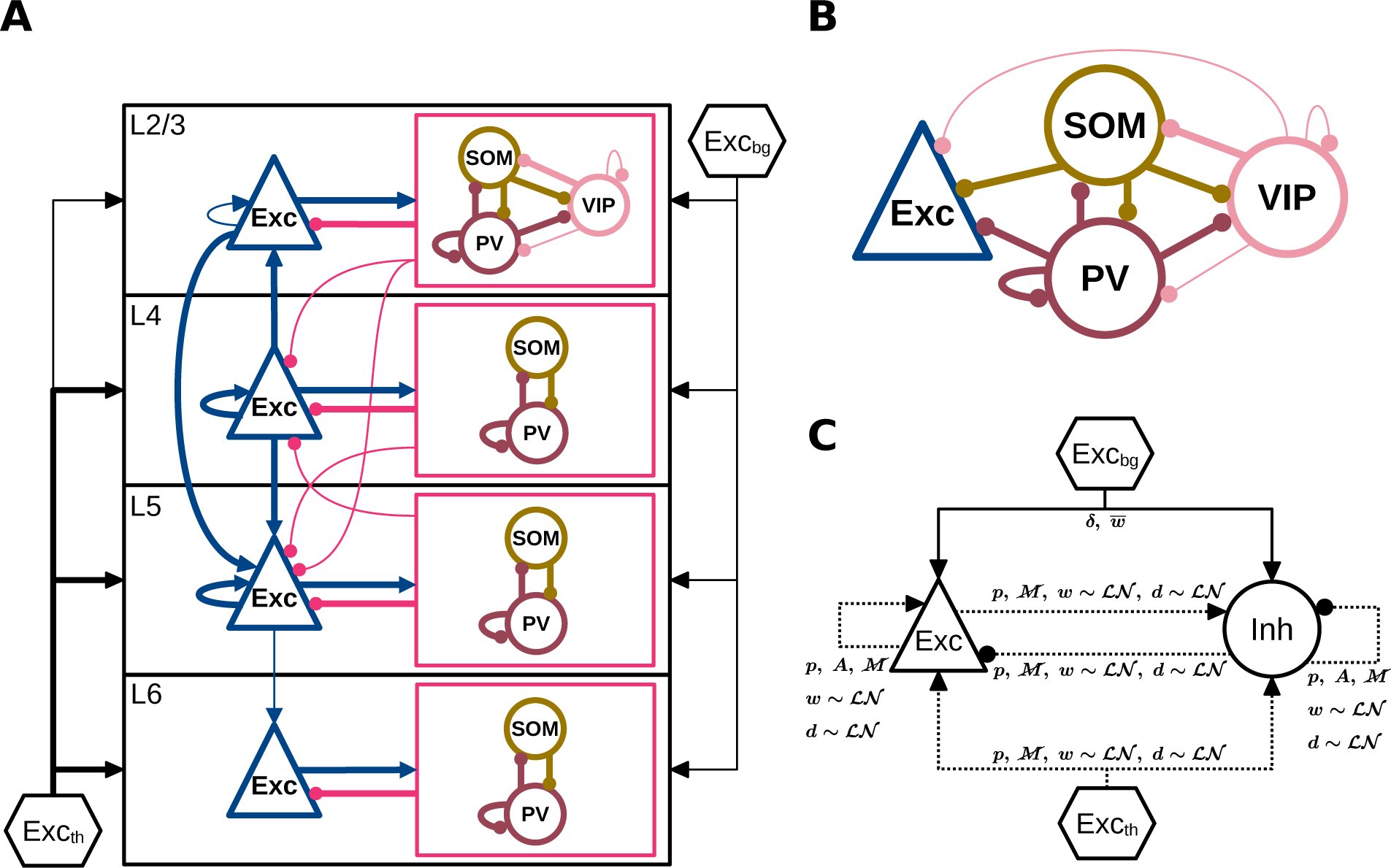
Model overview. (A) Populations and connectivity of the model. Exc: excitatory neurons. PV, SOM, VIP: parvalbumin-, somatostatin-, and vasoactive intestinal peptide-expressing inhibitory interneurons. Exc_bg_: background input. Exc_th_: thalamic input. For details of Exc_bg_ and Exc_th_, see Background Input and Thalamic Input, Methods. Interneurons in each layer are grouped by boxes (in magenta) to show their average external connections (lines to and from the box) and specific internal connections (lines within the box). Thin and thick lines show projections with a connection probability of 4–8% and ≥ 8% respectively. Those of *<* 4% are not shown. Note that in some cases this diagram may only partly reflect the true connectivity, due to limited availability of experimental data (see Model Parameters, Discussion). (B) L2/3 interneurons and their projections as an example of the cell-type-specific connectivity associated with interneurons. For simplicity, the excitatory projections are not shown. (C) Connection rules for excitatory (Exc) and inhibitory (Inh) populations regardless of layer and interneuron type, following the graphical notation introduced in Senk et al. (2022). The three interneuron types are considered together as Inh in (C). Symbol definitions for (C) are as follows. Solid and dashed lines: deterministic and probabilistic connections. *δ*: one-to-one connectivity. *p*: pairwise Bernoulli connectivity. *A*: autapses allowed. 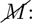 multapses not allowed. *w ∼ LN*: log-normally distributed synaptic weights. 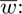 fixed synaptic weight. *d ∼ LN*: log-normally distributed synaptic delays.

### Model Components

#### Populations and Neuron Parameters

The model includes four cortical layers with Exc, PV, SOM, and VIP cells (Figure 1, Table 1). The cell numbers of populations in each layer are determined as follows. The layer-specific excitatory and inhibitory cell numbers (*N*_Exc_*, N*_Inh_) are specified according to estimates for the mouse C2 barrel column by Lefort et al. (2009). The PV, SOM, and VIP cell numbers (*N*_PV_*, N*_SOM_*, N*_VIP_) in each layer are determined by distributing *N*_Inh_:

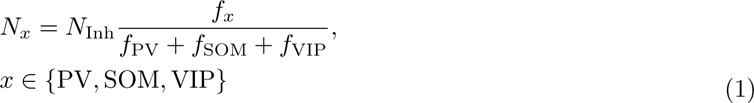

**Table 1.** Neural populations. The cell numbers are based on estimated excitatory and inhibitory neuron numbers from Lefort et al. (2009) and the relative quantities of PV, SOM, and VIP cells from Lee et al. (2010). For simplicity, all VIP cells are moved to L2/3. The numbers in parentheses show the VIP cell numbers before combining them into a single population.

| Layer | Exc | PV | SOM | VIP |
| --- | --- | --- | --- | --- |
| L2/3 | 1691 | 90 | 74 | 85 (67) |
| L4 | 1656 | 85 | 48 | (7) |
| L5 | 1095 | 109 | 105 | (7) |
| L6 | 1288 | 56 | 66 | (4) |

where *f*_PV_*, f*_SOM_*, f*_VIP_ are relative quantities of the three interneuron types, obtained from Figure 2D and 4D in Lee et al. (2010) with digitizing tools (see Relative Quantities of Interneuron Types, Supplementary Material). Since VIP cells are mostly located in L2/3 (Prönneke et al., 2015; Tremblay et al., 2016) and experimental data pertaining to their connectivity in other layers is lacking, we include all VIP cells in a single population for simplicity. This results in 13 populations in total (Table 1).

**Figure 2.**
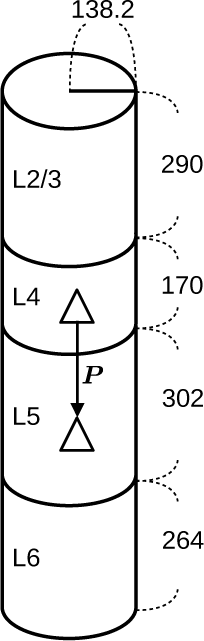
Dimensions of the microcircuit model in *µ*m. These dimensions are based on the data of a mouse C2 barrel column. The surface area is equivalent to 200*×*300 *µ*m (Petersen, 2019). The layer thicknesses are according to Lefort et al. (2009). Experimentally observed connection probabilities (*P*_exp_) are adjusted to derive model connection probabilities (*P*, see Figure 3) that correspond to these dimensions (see Derivation of Connection Probabilities, Supplementary Material).

The cell-type-specific membrane parameters (Table 2) are based on the *in vitro* data by Neske et al. (2015). Their L2/3 data are used for L2/3 and L4 of our model, and their L5 data for L5 and L6. Their L5 data include two excitatory subtypes, and we use weighted mean parameters of the two subtypes according to their relative cell numbers. The membrane time constants (*τ*_m_) are further adjusted to approximate the *in vivo* awake state as follows: Watson et al. (2008) reported a decrease in membrane resistance (*R*_m_) by 50.9% for excitatory and by 4.9% for inhibitory neurons from the Down state to the Up state. The Down and Up states are described as periods during which neuronal populations are silent and periods of long-duration multineuronal depolarization, respectively (Watson et al., 2008). We apply these reductions in *R*_m_ to the *τ*_m_ of the neurons in our model (Table 2), to approximate a change from a silent *in vitro* to the *in vivo* awake state.

**Table 2.**
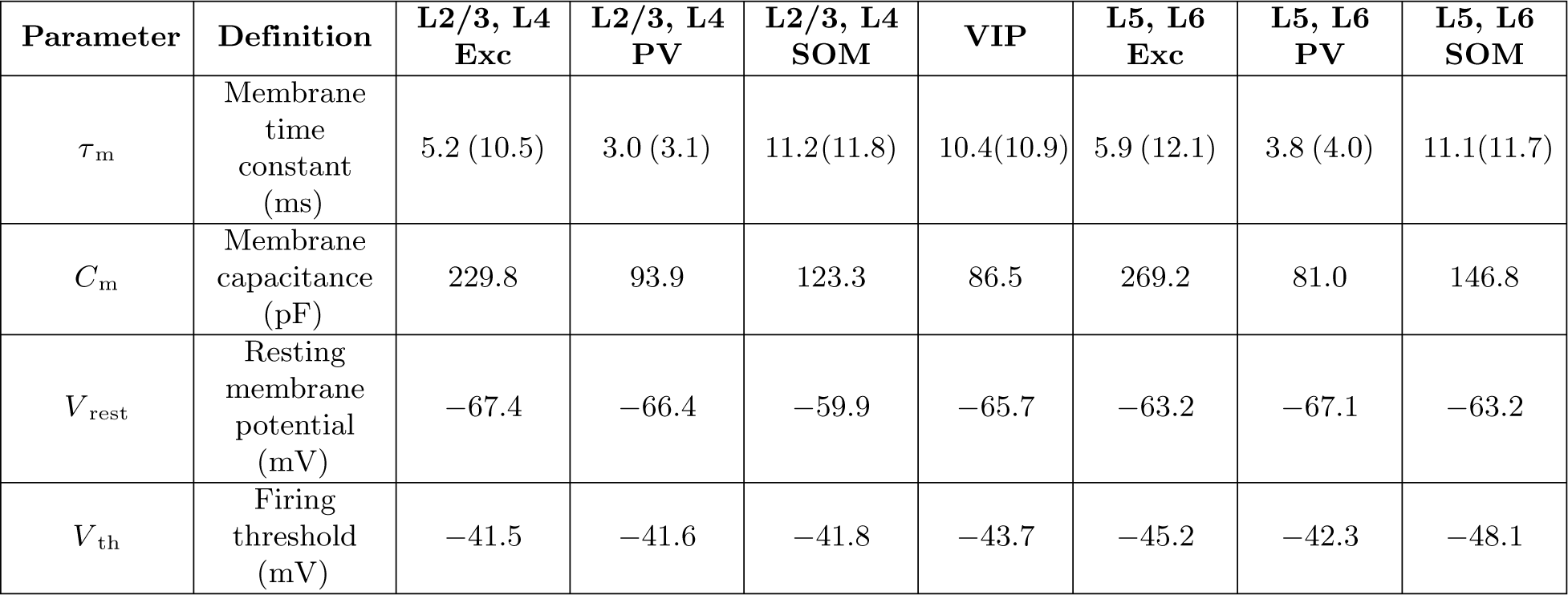
Layer- and cell-type-specific neuron parameters. The parameters are based on the data of supra- (L2/3) and infra-granular (L5) layers from Neske et al. (2015). The *τ* _m_ are adjusted to *in vivo* conditions according to the Down-to-Up-state decreases in membrane resistance of excitatory (50.9 %) and inhibitory (4.9 %) neurons reported by Watson et al. (2008) (see Populations and Neuron Parameters, Methods). Numbers in parentheses show *τ* _m_ before adjustment.

The absolute refractory period (*τ*_ref_) is 2.0 ms for every neuron, as in Potjans and Diesmann (2014).

#### Synaptic Parameters

Synaptic transmission events are modeled as currents with an instantaneous rise and a monoexponential decay. Cortical EPSPs consist of mainly the AMPA and the NMDA receptor-mediated components (Feldmeyer et al., 2002); however, only the AMPA component is considered here. This may be a reasonable approximation since, during ongoing activity, many NMDA receptors are presumably blocked due to their voltage dependence *in vivo* (Mayer et al., 1984). The synaptic weights (*w*) are determined as follows. First, the amplitudes of postsynaptic potentials (PSPs) are defined (Table 3). EPSPs of intracortical connections are set to 0.5 mV, which is consistent with the range of *in vivo* unitary EPSPs (Schoonover et al., 2014; Jouhanneau et al., 2015; Pala and Petersen, 2015; Jouhanneau et al., 2018). IPSPs are set to 2.0 mV, four times as strong as the EPSPs, as in Potjans and Diesmann (2014). EPSPs of the thalamic input follow the values of thalamocortical connections *in vivo* reported by Bruno and Sakmann (2006). For thalamic input to SOM cells, the PSPs are set to 50% of the others to reflect the reported weaker thalamocortical connections to this group of neurons (Ji et al., 2016). The intracortical EPSPs and IPSPs and thalamic EPSPs are log-normally distributed in the model to be consistent with *in vivo* (Schoonover et al., 2014; Jouhanneau et al., 2015; Pala and Petersen, 2015; Jouhanneau et al., 2018) and *in vitro* (Song et al., 2005) data. For intracortical EPSPs and IPSPs, the standard deviations are set to the same magnitude as the means (e.g., 0.5*±*0.5 mV). This is also consistent with the *in vivo* data (Schoonover et al., 2014; Jouhanneau et al., 2015; Pala and Petersen, 2015; Jouhanneau et al., 2018), where the standard deviations are 62 to 172% of the magnitude of the means. EPSP amplitudes of the background inputs are fixed to a value of 0.5 mV.

**Table 3.** Postsynaptic potentials. These PSP amplitudes are defined for the model with static synapses. The PSCs (Figure S1, left) required to produce such PSPs are calculated with Equation (2). In the model with STP, the PSPs and PSCs are dynamic, except for the background input.

| Parameter | Value (mean±SD or fixed) |
| --- | --- |
| EPSPs of intracortical connections | $0.5 \pm 0.5$ mV |
| IPSPs of intracortical connections | $-2.0 \pm 2.0$ mV |
| EPSPs of the thalamic input to Exc and PV | $0.49 \pm 0.13$ mV |
| EPSPs of the thalamic input to SOM | $0.245 \pm 0.065$ mV |
| EPSPs of the background input | 0.5 mV |

Subsequently, the amplitudes of exponentially decaying postsynaptic currents (PSCs) required to produce the defined PSPs in the different neuron types are calculated (Rotter and Diesmann, 1999; Maksimov et al., 2018):

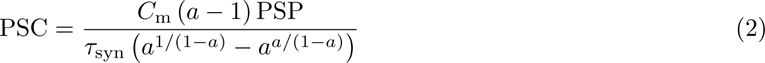

where *C*_m_ is the membrane capacitance, *τ*_syn_ is the PSC decay time constant, and *a* stands for 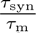 with *τ*_m_ the membrane time constant. *C*_m_ and *τ*_m_ depend on the postsynaptic population (Table 2), and *τ*_syn_ is 2 ms for excitatory (*τ*_syn,Exc_) and 4 ms for inhibitory (*τ*_syn,Inh_) connections (Table 4), which are chosen to approximate *in vitro* data from rat (Feldmeyer et al., 2002) and mouse (Ma et al., 2012), respectively. The resulting PSCs are shown in Figure S1A and are used as the synaptic weights in the model code.

**Table 4.** Synaptic time constants and delays. The time constants^1^ are chosen to approximate *in vitro* data of rats and mice (Feldmeyer et al., 2002; Ma et al., 2012). The delays^2^ are according to *in vivo* data of rats and mice (Bruno and Sakmann, 2006; Jouhanneau et al., 2018). For *d*_Exc_ and *d*_Inh_, values of pyramidal and PV cells in Jouhanneau et al. (2018) are used, respectively.

| Parameter | Definition | Value (mean±SD or fixed) |
| --- | --- | --- |
| $\tau_{\text{syn,Exc}}$ | Decay time constant of excitatory postsynaptic current | 2 ms |
| $\tau_{\text{syn,Inh}}$ | Decay time constant of inhibitory postsynaptic current | 4 ms |
| $d_{\text{Exc}}$ | Synaptic delay of recurrent excitatory connections | $1.36 \pm 0.51$ ms |
| $d_{\text{Inh}}$ | Synaptic delay of recurrent inhibitory connections | $1.43 \pm 1.09$ ms |
| $d_{\text{th}}$ | Synaptic delay of thalamic input | $1.72 \pm 0.73$ ms |
| $d_{\text{bg}}$ | Synaptic delay of background input | 0.1 ms |

The synaptic delays *d*_Exc_*, d*_Inh_*, d*_th_ (Table 4) follow *in vivo* data on excitatory and inhibitory intracortical connections (Jouhanneau et al., 2018) and thalamocortical connections (Bruno and Sakmann, 2006). They are log-normally distributed to be consistent with the experimental data as well. The synaptic delay of the Poisson background input (*d*_bg_) is fixed to the same value as the integration resolution (0.1 ms) because it is irrelevant for network activity.

#### Probabilities of Intracortical Connections

The intracortical connections are created with pairwise Bernoulli trials with projection-specific probabilities (*P*). Here, a projection stands for all connections from one neuron population to another (e.g., L4 E→L4 PV) (Senk et al., 2022). *P* of different projections are based on experimentally observed connection probabilities (*P*_exp_) in paired recordings of mouse barrel or somatosensory cortex *in vitro* (Galarreta and Hestrin, 2002; Kapfer et al., 2007; Lefort et al., 2009; Hu et al., 2011; Packer and Yuste, 2011; Ma et al., 2012; Xu et al., 2013; Pala and Petersen, 2015; Karnani et al., 2016b; Walker et al., 2016; Hilscher et al., 2017; Jouhanneau et al., 2018; Nigro et al., 2018; Scala et al., 2019). Since the spatial recording range differs among experiments, we adopt a spatial integration that uses *P*_exp_ to derive *P* corresponding to the model dimensions (Figure 2). The steps of this derivation are described in Derivation of Connection Probabilities, Supplementary Material. We use an exponential decay for the connection probability (Packer and Yuste, 2011; Perin et al., 2011), simplifying the distance dependence that is in reality shaped in more detail by factors including the geometry and density of the neurites. The pairwise Bernoulli rule creates only a single synapse between a pair of neurons, according to the given connection probability. Thus, it lumps together potentially multiple synapses (multapses; for terminology, see Senk et al. (2022)) forming each connection in the brain into a single synapse. By choosing larger synaptic weights to reflect potential multapses, the only dynamical difference that remains compared to modeling each synapse separately is that the corresponding transmission delays become identical. In return, the computational efficiency of the simulations is increased. Since paired recordings simultaneously activate all synapses of the source neuron onto the target neuron, the synaptic weights that we adopt to approximate unitary PSPs observed in such experiments (see Synaptic Parameters and Figure 3) naturally fit into this modeling scheme.

**Figure 3.**
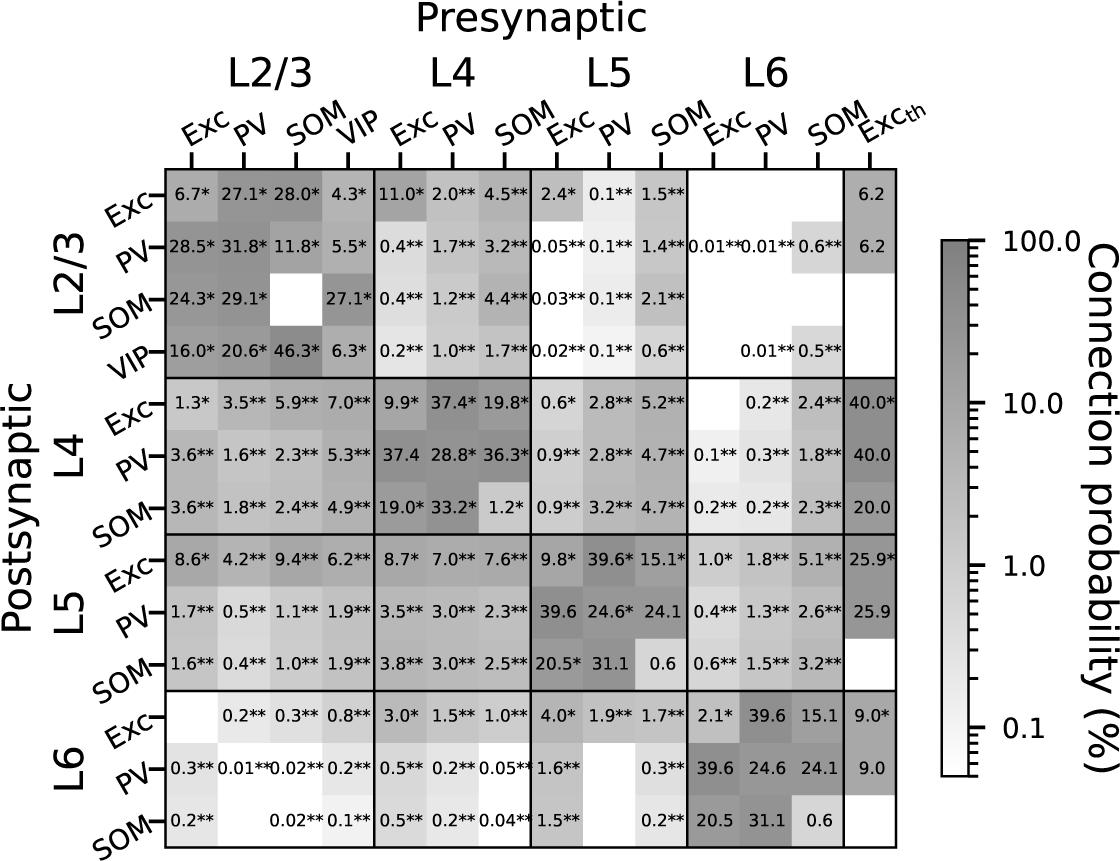
Connection probabilities (*P*) in %. *∗* indicates data derived from paired recording experiments. *∗∗* indicates estimations involving morphological data, based on Markram et al. (2015). Otherwise, data are based on assumptions. For the approaches to obtain these data, see Probabilities of Intracortical Connections, Methods, and Thalamic Input, Methods. Exc_th_: Thalamic input. Blanks: Probability is zero.

Where experimental data are unavailable, we use assumed or estimated values for connection probability (Figure 3). For the intra-layer projections, the following rules are applied. (1) Exc→PV equals PV→Exc. This is based on the experimental observation that connections between pairs of Exc and PV cells have a very high probability of reciprocation (Geiger et al., 1997; Koelbl et al., 2015; Couey et al., 2013; Hu et al., 2014; Qi et al., 2017). (2) L5 Exc→SOM, SOM→Exc and SOM→SOM use the averages of L2/3 and L4, due to the lack of more precise constraints. (3) For L6 Exc→Exc, the connection probability is derived from experimental data (Lefort et al., 2009). The other projections in L6 use the same values as their counterparts in L5, also due to the lack of more precise constraints. For most inter-layer projections, physiologically determined connection probabilities are lacking. Therefore, we use the estimated connection probabilities from Markram et al. (2015)^2^ as the *P*_exp_ to derive *P*. Since their algorithm is based on morphological data, we map their morphological types to our cell types as follows. Following Figure 2 and Table 1 in Markram et al. (2015), we take the pyramidal cells, star pyramidal cells, and spiny stellate cells as Exc cells, the large basket cells and normal basket cells as PV cells, the Martinotti cells as SOM cells, and the double bouquet cells and bipolar cells as VIP cells. Accordingly, for each layer the connection probabilities of morphological subtypes are combined by weighted averaging, taking into account their cell numbers in the database, to obtain the connection probabilities for Exc, PV, SOM, and VIP cells.

Overall, 29 of 43 intra-layer connection probabilities and 12 of 126 inter-layer connection probabilities are directly derived from experimental data. For the others, it is still necessary to use assumptions and estimations as described above.

#### Background Input

The background input (Exc_bg_) for each cell is a homogeneous Poisson spike input with a fixed EPSP amplitude of 0.5 mV and a constant but cell-type-specific firing rate (*r*_bg_). The *r*_bg_ for each cell type is optimized to obtain plausible population firing rates (see Parameter Optimization and Model Simulations in the following). The background input is always present in all simulations in this study.

#### Thalamic Input

To model a thalamic input (Exc_th_), we estimate the cell number of a barreloid in the ventral posteromedial (VPM) nucleus of mouse corresponding to the C2 whisker. We derive a number of 115, by dividing the total cortical neuron number of our model (=6448) by the “Ratio S1/VPM” of rat C2 (=56) from Table 1 in Meyer et al. (2013). However, with firings according to touch-evoked responses of VPM (Figure 4), this number of cells produces much smaller cortical responses (data not shown) in our model than *in vivo* (Yu et al., 2019). Therefore, we double it to create an Exc_th_ of 230 cells for larger cortical responses. We consider this adjustment biologically plausible, as neurons in the barrel cortex can respond to several adjacent whiskers with similar response latencies, i.e., they have multi-whisker receptive fields (Bruno and Simons, 2002; Le Cam et al., 2011; Staiger and Petersen, 2021).

**Figure 4.**
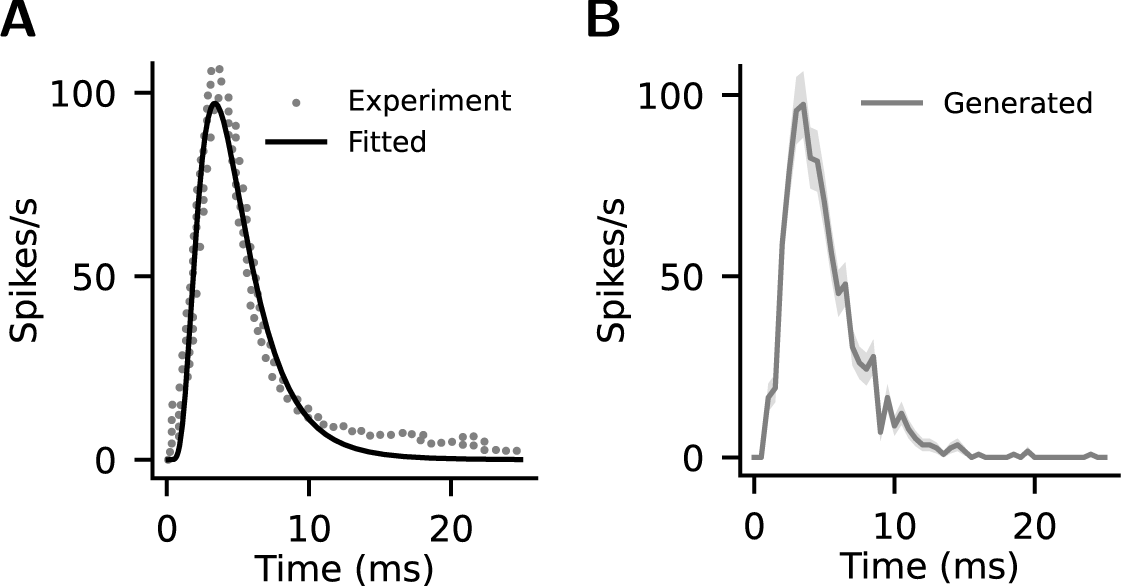
Evolution of thalamic (VPM) firing rate in response to whisker touch. (A) Experimental data digitized from Figure 3F in Yu et al. (2019) (gray dots) and the time course fitted with a log-normal function (black line). This fitted time course is used for the simulated thalamic input (Exc_th_) (see Thalamic Input, Methods). (B) Mean firing rates actually generated in the thalamic neurons (n=230) in one simulation instance. Shaded area: SEM.

The Exc_th_ is connected to Exc and PV cells in all layers. SOM cells are targeted only in L4, and VIP cells (located only in L2/3 in our model) are not targeted. This arrangement follows the pattern of cell-type-specific thalamocortical projections reported by Ji et al. (2016). The layer-specific connection probabilities (Figure 3) are specified according to the *in vivo* data from Constantinople and Bruno (2013). However, they did not report the connection probability for L2/3. Therefore, we estimate it as follows. We calculate the L2/3-to-L4 ratio 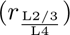 of the thalamocortical synapse number per cortical neuron according to the data from Oberlaender et al. (2012). Then, assuming the thalamocortical connection probability is proportional to the average thalamocortical synapse number per neuron (i.e., assuming the same multiplicity of synapses per pair of cortical and thalamic cells for the two layers), the L4 connection probability in Constantinople and Bruno (2013) is multiplied by 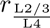 to obtain the estimated L2/3 connection probability (Figure 3, Exc_th_→L2/3 Exc and Exc_th_→L2/3 PV). The synaptic weights are specified according to the *in vivo* data from Bruno and Sakmann (2006) (Table 3). We use these probabilities and weights for Exc and PV cells. For SOM cells, we use 50% of these values (Figure 3, Table 3) to reflect their smaller innervation percentages and strengths reported by Ji et al. (2016).

The thalamic stimulation is generated according to the touch-evoked responses of VPM from Figure 3F in Yu et al. (2019). The data are obtained with digitizing tools and offset along the y-axis to zero at stimulus onset (Figure 4A, gray dots), and then fitted with a log-normal function to produce a firing rate time course with 0.1 ms resolution (Figure 4A, black line). This time course is generated in the form of inhomogeneous Poisson spike trains in all thalamic neurons to serve as a transient thalamic input for the model. The Poisson spike train of each neuron is generated according to this time course but individually randomized by NEST, hence producing a certain variability. Figure 4 shows the time course of the mean firing rate that is actually generated in the 230 thalamic neurons in one simulation instance (Figure 4B, gray line).

#### Short-Term Synaptic Plasticity

We created another version of the model that includes all components described in the preceding but further incorporates STP synapses. The STP synapses are implemented as in the Tsodyks model (Tsodyks et al., 2000). This includes a parameter determining the dynamics of synaptic release probability (U), a facilitation time constant (F), and a depression time constant (D).

Values of U, F, and D for different projections are fitted using experimental unitary PSPs that demonstrate STP (Kapfer et al., 2007; Ma et al., 2012; Hu et al., 2014; Lefort and Petersen, 2017). For each projection (e.g., L4 Exc→L2/3 Exc), we created a pair of neurons connected with an STP synapse. Then, we let the presynaptic neuron fire at the same constant rate as in the corresponding experiment. With this pair of neurons, we ran repeated simulations scanning U, F, and D (U: 0.05–1.0 in steps of 0.05; F, D: 0–1000 ms in steps of 20 ms) to determine the best-fit parameters, i.e., the set of U, F, and D that yields the smallest root-mean-square error (RMSE) of normalized PSP amplitudes between simulation and experiment (see Figure 5A for examples). The resulting set of U, F, and D is used for this particular projection. Note that Lefort and Petersen (2017) and Ma et al. (2012) subtracted overlapping components of preceding PSPs before calculating PSP amplitudes. Such a subtraction is also performed for the corresponding simulation data to ensure an accurate fit (e.g., L4 Exc→L2/3 Exc and L5 Exc→L6 Exc in Figure 5A).

**Figure 5.**
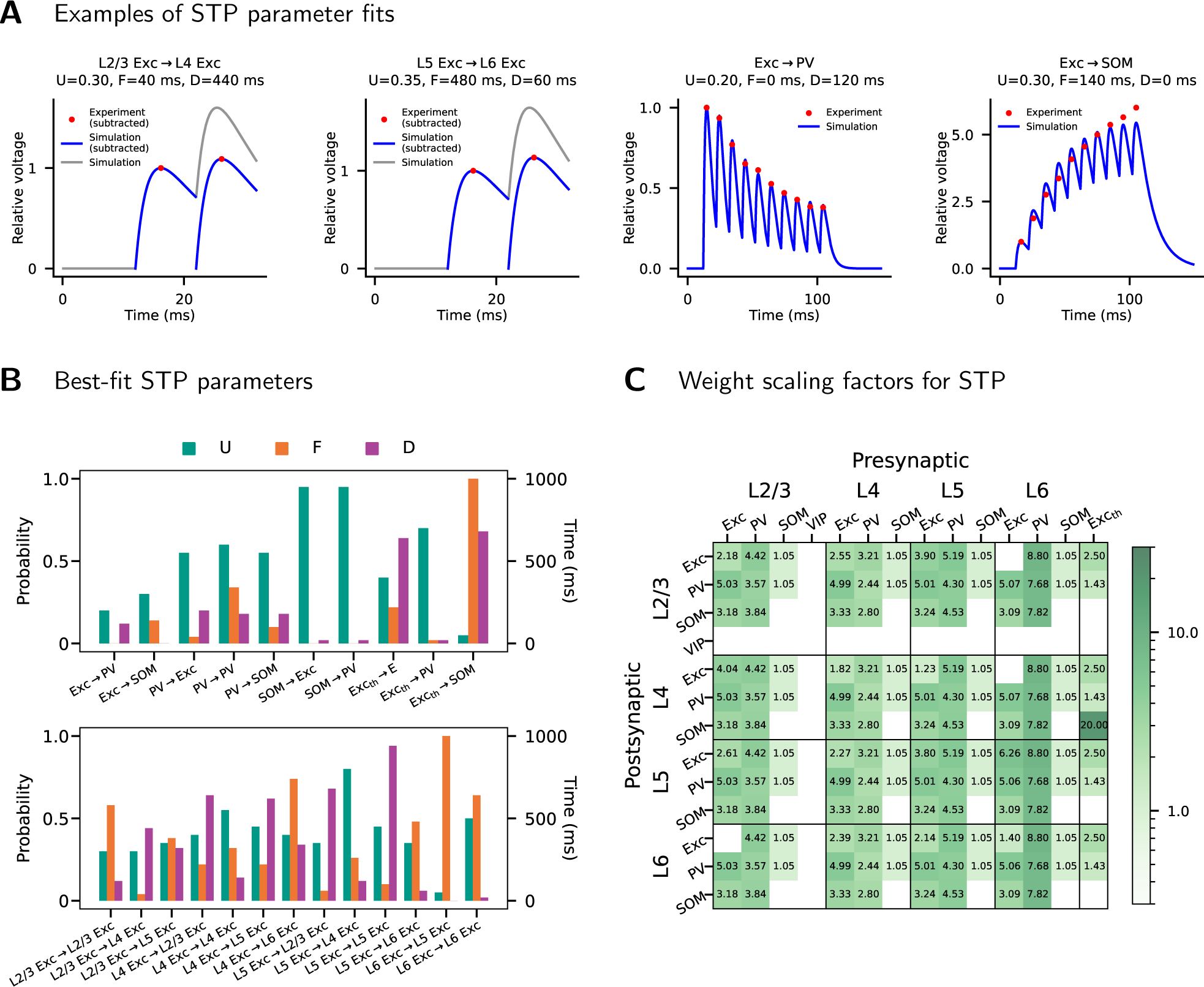
Short-term plasticity parameters. (A) Examples of STP parameter fits for four projections. U, F, and D are STP parameters. U: parameter for release probability. F: facilitation time constant. D: depression time constant. Blue lines: Simulated PSPs with the best-fit STP parameters (the values shown above). Red dots: Experimental data on PSP amplitudes. In all cases, PSP data are normalized to the amplitude of the first PSP. For each projection, the best-fit STP parameters are determined by minimizing the RMSE of the PSP amplitudes between simulation and experiment. If the experimental data include a subtraction of preceding PSPs, such a subtraction is also done for the simulated PSPs, e.g., L4 Exc→L2/3 Exc and L5 Exc→L6 Exc (gray lines show the simulated PSPs before subtraction). (B) Best-fit STP parameters of all projections with STP. These best-fit STP parameters are used throughout this study for the model with STP. Projections that remain static are not shown. Exc_th_: thalamic input. (C) Mean weight scaling factors across 20 simulation instances for the projections with STP. In each simulation, the weight scaling is used to yield a resting state approximating the model with static synapses (see Short-Term Synaptic Plasticity, Methods). Blanks: Connection probability is zero or the synaptic weight is static.

With STP incorporated, the synaptic weights change during simulation. Therefore, we scale the synaptic weights of the projections with STP, so that they evolve to a state that approximates the defined static synaptic weights (PSPs as in Table 3, PSCs as in Figure S1). This results in a model with the fitted STP parameters and population firing rates similar to the original model, ensuring close comparability between the two model versions. To obtain the scaling factors that achieve this, we used a numerical approach as follows. For each projection, we created a pair of neurons connected with its fitted STP parameters. Then, we let the presynaptic neuron fire at the same rate as the corresponding population in the model with static synapses. With this pair of neurons, we ran repeated 5-s simulations while scaling the initial synaptic weight, until the last synaptic weight recorded from the simulation deviated less than 0.1 pA from the target (i.e., the corresponding PSC in Figure S1A). The weight scaling factor thus obtained is used for this particular projection. Note that because this approach relies on the simulated population firing rates in the model with static synapses, the obtained scaling factors always depend on the random seed and the background inputs used for simulation. Figure 5C shows the mean scaling factors across 20 simulation instances determined with this approach.

For the two fitting processes described above, the membrane parameters of the postsynaptic neuron are set to those of the corresponding population (as in Table 2). In the fitting for weight scaling, we use presynaptic spike trains with fixed inter-spike intervals, as this turned out to yield population firing rates that better approximate the original model than fitting with Poisson statistics or spike trains taken directly from the original model.

For the thalamic input, the STP parameters are derived from the data by Hu et al. (2014) with the same approach described above. However, because the thalamic input itself is transient, the described approach for weight scaling does not apply. Instead, the weight is simply scaled such that the effective initial weight equals the originally defined weight *w*. Assuming the scaled weight is *w^′^*, then the effective initial weight is *u × w^′^*, where *u* is the initial synaptic release probability (Tsodyks et al., 2000). Therefore, we set *w^′^* to 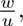 yielding effective initial weight 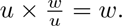

### Parameter Optimization and Model Simulations

In Results, we first present the simulations for the resting state (Modeled Resting State, Results) and cell-type-specific stimulation (Network Responses to Cell-Type-Specific Stimulation, Results). The model for this part is optimized with respect to only the firing rates of the background inputs (*r*_bg_). This optimization is cell-type-specific but layer-independent to limit the number of fitted parameters and thereby increase the robustness of the model. Different values of *r*_bg_ for Exc, PV, SOM, and VIP cells are scanned to find a combination that results in the smallest deviation in population firing rates from the *in vivo* data (Yu et al. (2019); See Table 5 for the criteria values). This only includes L2/3 and L4 because incorporating L5 and L6 results in large deviations (data not shown), possibly due to insufficient experiment-based connection probability data and consequent lesser reproduction of the *in vivo* activity. The deviation is quantified by the root-mean-square percentage error of the mean population firing rates (*t* = 10 s to *t* = 15 s) across simulation instances, i.e., for each population, the difference between the simulated and the *in vivo* data is computed as a fraction of the *in vivo* value, and the root mean square of the resultant values is calculated. The parameter scan for this optimization is done in two steps, first using a coarse interval (500 spikes/s) and then a finer one (100 spikes/s). The resultant best-fit *r*_bg_ combination is used throughout this study unless otherwise specified. This optimization process (illustrated in Optimization of the Background Input, Supplementary Material) is done separately for the cases with static synapses and with STP. The resultant model versions are named Base and Base-STP respectively.

**Table 5.** Neuronal firing rates of the optimized model. The values represent mean *±* SD, median, 25th–75th percentile, and cell number, for the model with static synapses (Base), the model with STP (Base-STP), and the experimental data (Experiment). For Base and Base-STP, the cell numbers represent all neurons in 20 randomized simulations (n = 20 *×* population cell number in Table 1). The experimental data are from the non-whisking-state firing rates in Table 1 of Yu et al. (2019). The mean firing rates in L2/3 and L4 (indicated by *) are used for background input optimization (see Parameter Optimization and Model Simulations, Methods).

|  |  | Exc | PV | SOM | VIP |
| --- | --- | --- | --- | --- | --- |
| L2/3 | Base | 2.1 ± 2.0, 1.6, 0.8-2.8, n = 33820 | 15.6 ± 7.6, 14.6, 9.8-20.2, n = 1800 | 2.5 ± 3.2, 1.4, 0.4-3.2, n = 1480 | 10.2 ± 7.1, 8.6, 4.6-14.4, n = 1700 |
|  | Base-STP | 2.2 ± 1.8, 1.8, 1-3, n = 33820 | 16.2 ± 6.9, 15.2, 11-20.6, n = 1800 | 2.7 ± 3.0, 1.6, 0.6-3.6, n = 1480 | 13.3 ± 8.2, 12, 7-18.6, n = 1700 |
|  | Experiment | 2.7* ± 3.7, 0.6, 0.5-4.5, n = 5 | 13.8* ± 8.9, 11.7, 7.5-23.3, n = 8 | 2.6* ± 3.6, 0.4, 0.03-4.1, n = 9 | 14.6* ± 7.3, 11.1, 8.5-21.0, n = 9 |
| L4 | Base | 0.5 ± 0.6, 0.4, 0.0-0.8, n = 33120 | 10.4 ± 5.3, 9.6, 6.4-13.6, n = 1700 | 2.5 ± 2.8, 1.6, 0.6-3.6, n = 960 | - |
|  | Base-STP | 0.6 ± 0.6, 0.4, 0.2-0.8, n = 33120 | 10.5 ± 4.8, 9.6, 6.8-13.4, n = 1700 | 2.9 ± 2.9, 2, 0.8-4.2, n = 960 | - |
|  | Experiment | 0.5* ± 0.8, 0.0-0.7, n = 95 | 10.2* ± 7.2, 7.8, 4.3-14.7, n = 43 | 2.6* ± 3.2, 0.6, 0.3-4.9, n = 27 | - |
| L5 | Base | 2.0 ± 3.1, 0.8, 0.2-2.6, n = 21900 | 20.2 ± 12.1, 18.4, 11-27.3, n = 2180 | 1.0 ± 2.1, 0.2, 0.0-0.8, n = 2100 | - |
|  | Base-STP | 2.3 ± 3.0, 1.2, 0.4-3, n = 21900 | 21.0 ± 10.1, 19.6, 13.6-27, n = 2180 | 2.7 ± 3.9, 1.2, 0.2-3.6, n = 2100 | - |
|  | Experiment | 6.8 ± 5.2, 5.2, 2.7-11.2, n = 23 | 7.5 ± 5.2, 7.6, 4.3-8.7, n = 7 | 2.8 ± 4.5, 0.8, 0.2-3.6, n = 18 | - |
| L6 | Base | 3.0 ± 5.5, 0.8, 0.0-3.2, n = 25760 | 38.4 ± 21.2, 36.6, 22.0-53.5, n = 1120 | 5.8 ± 9.2, 1.8, 0.2-7.6, n = 1320 | - |
|  | Base-STP | 3.9 ± 5.2, 2, 0.6-5.2, n = 25760 | 33.2 ± 15.7, 32.2, 21.0-44.2, n = 1120 | 23.7 ± 16.3, 20.8, 11.2-33.4, n = 1320 | - |
|  | Experiment | 6.1 ± 6.9, 2.6, 0.4-11.5, n = 30 | 16.9 ± 14.3, 17.2, 4.6-22.0, n = 15 | 3.9 ± 4.9, 1.7, 0.5-6.9, n = 26 | - |

Subsequently, we compare the thalamic-input-evoked cortical responses of the model with the experimentally observed touch-evoked responses from Yu et al. (2019) (Network Responses to Thalamic Stimulation, Results). In this part, we start from the Base model and scan several selected model parameters while evaluating the evoked responses. The result of this scan demonstrates how the selected parameters contribute to the reproduction of the experimentally observed responses. The model with the best-fit responses in this part is named the thalamocortical-adjusted (TC-adjusted) model. The corresponding version with STP (TC-adjusted-STP) is created by applying the same parameter changes to the Base-STP model.

In the following, we describe the methods used for each of the simulation results presented in this study.

#### Simulation of Network Resting State

The resting state of our model is simulated using only the background (homogeneous Poisson) input. The neuronal firing rates and the asynchronous irregular (AI) activity of this state are calculated and compared with experimental criteria. The experimental firing rates are taken from Yu et al. (2019). The AI activity is evaluated according to the study by Maksimov et al. (2018). AI activity corresponds to a state of low synchrony and irregular neuronal spiking, which exists in LIF neuron networks with sparse connectivity and balanced inputs (Brunel, 2000) as well as *in vitro* and *in vivo* samples (Steriade et al., 2001; Destexhe et al., 2007; Dehghani et al., 2016). Based on *in vivo* data, Maksimov et al. (2018) proposed criteria on cortical AI states for modeling studies. The study computes the pairwise spike count correlation and the coefficient of variation of inter-spike intervals (CV ISI) for *in vivo* data from rat and macaque (Chu et al., 2014; Watson et al., 2016). While the values were largely similar for the two species, we here consider the criteria based on rat data, which were obtained by Maksimov et al. (2018) as follows: For both correlation and CV ISI, means of each of 13 recording sessions from awake rat frontal cortex were calculated. For CV ISI, the mean across Up states in anesthetized rat motor cortex is further provided.

We evaluate the AI activity of our model with an approach akin to that of Maksimov et al. (2018). For each layer, 200 neurons are randomly chosen regardless of cell type, to calculate the mean pairwise spike count correlation. The same number of neurons are chosen separately to calculate the CV ISI. Data from *t* = 10 s to *t* = 15 s of the simulations are used. For pairwise spike count correlation, the bin width is 10 ms, and neurons with no spikes during the sampled period are excluded before selection. For CV ISI, neurons with a firing rate lower than 1 spikes/s during the sampled period are excluded before selection. The results are compared with the awake and anesthetized state criteria for rat established by Maksimov et al. (2018).

#### Simulations of Cell-Type-Specific Stimulation

To study how the network responds to activation of different cell types, an additional homogeneous Poisson input with a fixed EPSP amplitude of 0.5 mV is applied to each cell type in L2/3 and L4. The duration and interval of this input are both 1 s, and it is repeated 20 times and for nine levels of firing rate (*r*_stim_). The levels include *r*_stim_ = 0 spikes/s and eight other levels, which depend on cell type and reach up to maximally 1000 spikes/s. The resultant population firing rates in the same layers are calculated for the later half of each stimulation period (*t* = 500 ms to *t* = 1000 ms after stimulus onset). This protocol is performed for Exc, SOM, PV, and VIP populations and for L2/3 and L4 separately in dedicated simulations.

We statistically test the stimulus effects on the population firing rates in the same layer. The population firing rates are first normalized to the case of no stimulation (*r*_stim_ = 0 spikes/s). The data for each stimulus level are then tested for significant deviation from 1 (two-tailed one-sample t-test, for 20 simulation instances).

#### Simulations of Thalamic Stimulation

The network responses to thalamic input (see Thalamic Input) are similarly studied with dedicated simulations. In each simulation, the thalamic stimulation is applied every 1 s and repeated 10 times. Responses of cortical neurons are evaluated in terms of population peristimulus time histogram (PSTH) in 0.5 ms time bins. For each population, amplitude (*a*_peak_) and time (*t*_peak_) of the peak in the PSTH from *t* = 0 to *t* = 50 after stimulus onset are calculated. For comparison, *a*_peak_ and *t*_peak_ are obtained from the PSTHs of touch-evoked cortical responses in Figure 3F of (Yu et al., 2019), which also includes the data up to *t* = 50 after stimulus onset. Similarity between simulation and experiment is quantified via the RMSEs of *a*_peak_ and *t*_peak_.

### Hardware and Software Configurations for Simulations

NEST 3.6 (Villamar et al., 2023)^3^ is used for model implementation and simulation. All simulations are executed on a computing cluster with 48-core compute nodes running at 2.5 GHz. The simulation resolution is 0.1 ms in all cases. Each simulation used 24 cores on one compute node, with OpenMP^4^ for parallel computing. For a resting-state simulation of 15 s biological time, both model versions take approximately 0.2–0.25 core-hours (*∼*2 s build phase and 30–35 s simulation phase on 24 cores).

For the parameter scans described in Parameter Optimization and Model Simulations, 10 simulation instances are run for each parameter setting. The other results each rely on 20 simulation instances. Each simulation instance uses a unique seed for the randomization of Poisson input and network connectivity.

The simulation duration (biological time) is 15 s for the resting state (Simulation of Network Resting State) and the optimization for *r*_bg_ (Parameter Optimization and Model Simulations), 370 s for the cell-type-specific stimulation (Simulations of Cell-Type-Specific Stimulation), and 20 s for all simulations with thalamic input (Parameter Optimization and Model Simulations, Thalamic Input). In all cases, the first 10 s of simulation is excluded from analysis to avoid transients.

## Results

### Parameter Fits for Short-Term Synaptic Plasticity

As described in Methods, we fit the Tsodyks model to experimental STP data. Figure 5B shows the fitted STP parameters. Projections not shown in the figure remain static. For Exc→Exc projections (Figure 5B, bottom), experimental STP data from different layers are taken from Lefort and Petersen (2017). For projections involving interneurons (Figure 5B, top), experimental STP data are limited to L2/3 and L4. Kapfer et al. (2007) reported data involving L2/3 pyramidal cells, fast-spiking (FS) cells (taken as PV cells in our model), and SOM cells (pyramidal→FS, pyramidal→SOM). Ma et al. (2012) reported data involving L4 excitatory regular-spiking (RS) cells, FS cells, and SOM cells (FS→RS, FS→FS, FS→SOM, SOM→RS, SOM→PV). Since, for each of these projections, data are available only from one layer (L2/3 or L4), for these projections we use the fitted parameters regardless of layer in our model.

Karnani et al. (2016b) also report experimental STP data involving different cortical interneuron types, including VIP cells. However, adding this dataset resulted in an overactive resting state of the model (data not shown); therefore, it is not incorporated in this study.

Hu and Agmon (2016) report experimental STP data on cell-type-specific thalamocortical projections. The averaged data on FS and SOM cells are listed in their Figure 1, but those on excitatory cells are not provided. Therefore, we digitized the data on FS and SOM cells (cell-attached data in the case of SOM cells; see their Figure 1R) for the STP of Exc_th_→PV and Exc_th_→SOM projections in our model. For Exc_th_→E, we use the STP of L4 Exc→L2/3 Exc in our model as a substitution (Figure 5B), which we likewise consider a feedforward projection in sensory processing.

Figure 5C shows the mean weight scaling factors across 20 simulation instances for the model version with STP. Figure S1 shows the resulting mean resting-state weights across 20 simulation instances compared to the model version with static synapses. No weight scaling is performed for projections that remain static.

### Modeled Resting State

The resting states of the optimized model versions with static synapses (Base) and with STP (Base-STP) are shown in Figure 6. The statistics of neuronal firing rates across layers and cell types are listed in Table 5 along with the data used for their optimization (Yu et al., 2019).

**Figure 6.**
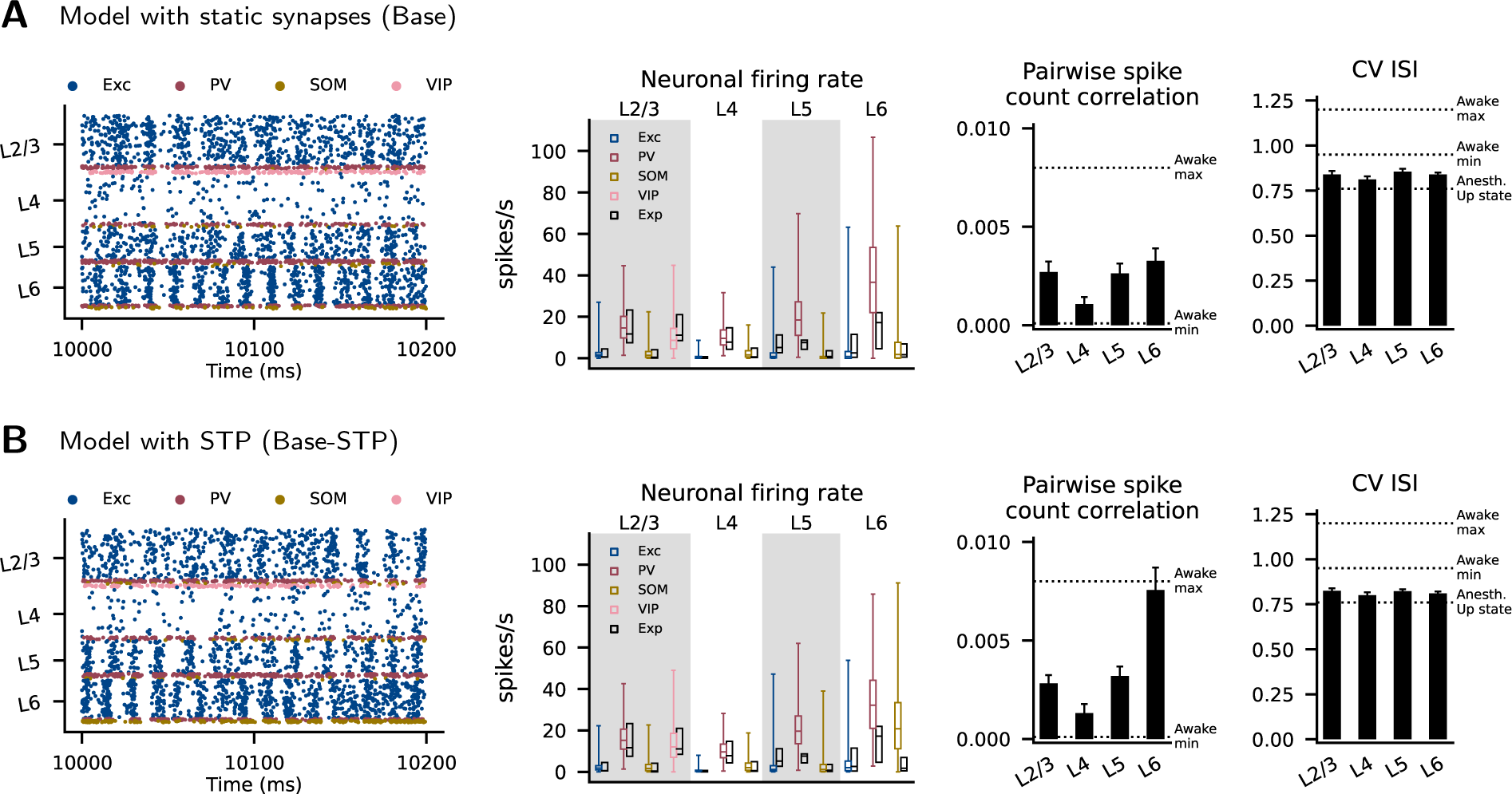
Resting state of the model with static synapses (A) and with STP (B). Left to right: raster plot of neuronal firings, neuronal firing rate of each cortical population, pairwise spike count correlation in each layer, and coefficients of variation of inter-spike intervals (CV ISI) in each layer. For the neuronal firing rate, the colored boxplots show the medians, 25th and 75th percentiles, and ranges for all neurons from 20 simulation instances (see Table 5 for numerical values). The black boxplots show the corresponding experimental data from Table 1 in Yu et al. (2019) (full ranges are not available). For the pairwise spike count correlation and CV ISI, the black bars show means and standard deviations across 20 simulations, and the dashed lines show the in vivo criteria as follows. Awake max, Awake min: the maximum and minimum of the means for 13 recording sessions in frontal cortices of awake rats. Anesth. Up state: the mean of recorded Up states in motor cortex neurons of anesthetized rats. These values were provided by Maksimov et al. (2018).

To evaluate the asynchronous irregular (AI) activity of the model, we use the *in vivo* criteria on pairwise spike count correlation and coefficient of variation of the inter-spike intervals (CV ISI) established by Maksimov et al. (2018) (see Simulation of Network Resting State, Methods). We consider the criteria based on awake and anesthetized states in rats. For both model versions, the mean layer-specific correlations are always within the range of the awake states, and the mean layer-specific CV ISIs are always between the awake and anesthetized states (Figure 6). This shows that the modeled resting state has an AI activity similar to the *in vivo* condition. Nevertheless, the Base-STP model shows a correlation in L6 more than two times that in the Base model.

Overall, the differences between the activity statistics of the two model versions are small, indicating that STP has only a limited effect on ongoing activity. We also examined the resting states in several model versions with alternative parameter settings, giving the same qualitative results (Figure S4, S7, S10, and S11).

To confirm the roles of the recurrent and background inputs in the model, we perform resting-state simulations where certain recurrent connections are removed (Figure S17). While removing the interlaminar connections does not change the resting state very much, removing the intralaminar connections causes an overactive state where some Exc populations fire at *>* 100 spikes/s. This indicates that the intralaminar connections are important in maintaining the resting state. Thus, the simulated activity is shaped by both the recurrent and the background inputs. Since there is no inhibitory background input, the inhibitory interneurons provide the stabilizing force in maintaining the low-rate activity.

### Network Responses to Cell-Type-Specific Stimulation

To assess inhibitory and disinhibitory effects of the interneuron types, we simulated how cell-type-specific stimulation changes network activity in L2/3 and L4 (Figure 7 and 8). In L2/3 of the Base model, PV and SOM activation evoke inhibition while VIP activation evokes disinhibition: The Exc firing rate decreases significantly with PV (-46.4% at *r*_stim_ = 1000 spikes/s, *p <* 0.001) and SOM (-72.2% at *r*_stim_ = 200 spikes/s, *p <* 0.001) activation, while it increases significantly with VIP activation (+10.4% at *r*_stim_ = 200 spikes/s, *p* = 0.004). In L4, PV activation evokes inhibition while SOM activation evokes disinhibition: The Exc firing rate decreases significantly with PV activation (-84.8% at *r*_stim_ = 1000 spikes/s, *p <* 0.001), while it increases significantly with SOM activation (+11.0% at *r*_stim_ = 200 spikes/s, *p <* 0.001).

**Figure 7.**
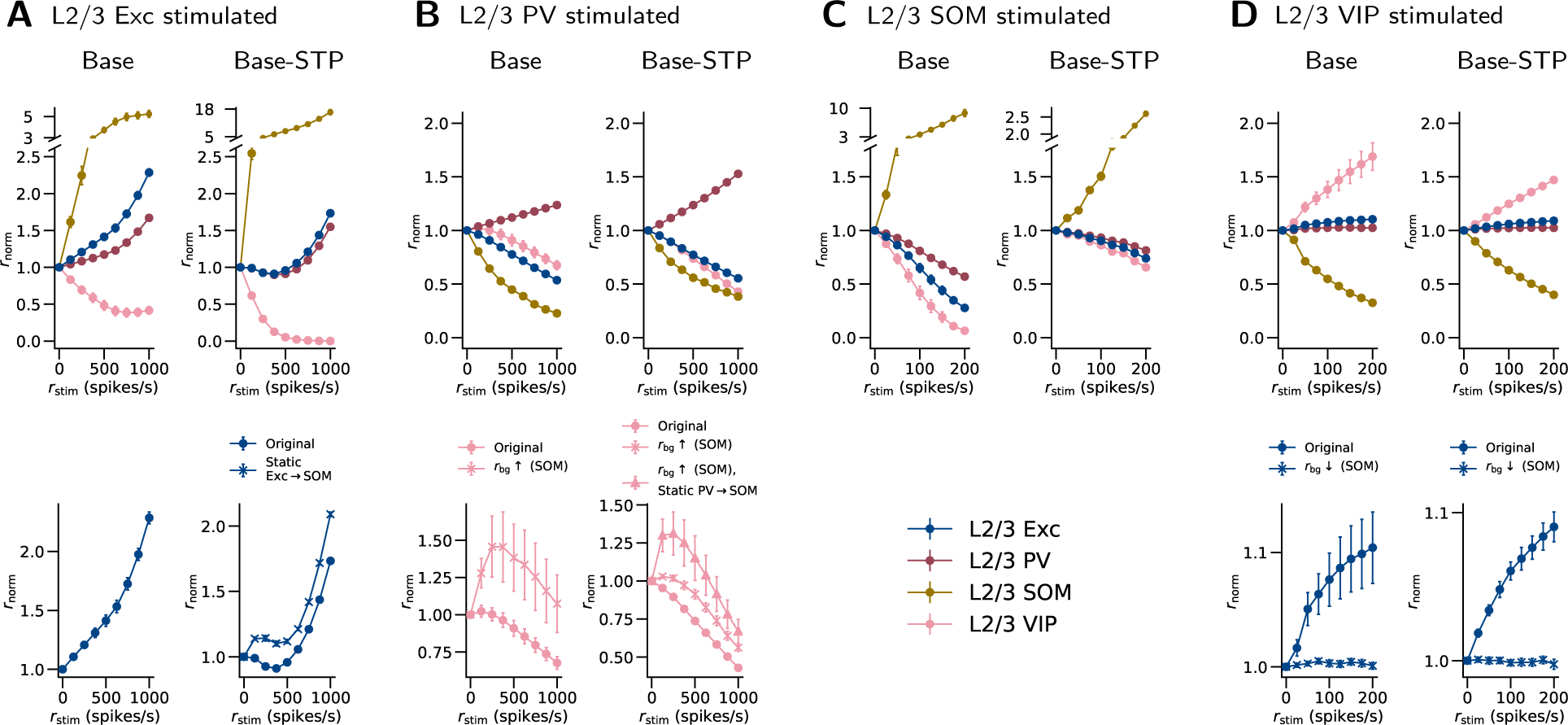
L2/3 network responses to cell-type-specific stimulation. Each of (A) to (D) shows the changes in population firing rates when one particular cell type is stimulated in L2/3. Through stimulating one cell type and observing the responses in the other cell types, these results demonstrate the roles of the different interneuron types. *r*_stim_: rate of stimulation applied. *r*_norm_: resulting relative population firing rate, normalized to the data at *r*_stim_ = 0. The first row shows results for all L2/3 populations in the original models (Base and Base-STP). The second row compares responses of a specific population between the original models and modified models, which reveals effects involving interneurons and their STP (see also Figure 9 for illustration). Original: original model. Static X→Y: STP in X→Y connections is excluded, X and Y being two cell types. *r*_bg_↑ (X), *r*_bg_↓ (X): *r*_bg_ for cell type X is increased or decreased. Details of the stimulation protocol are described in Simulations of Cell-Type-Specific Stimulation, Methods. Error bars: SEM across simulation instances (n=20) with different randomization seeds. In some cases the SEM is so small that the bar is not visible.

**Figure 8.**
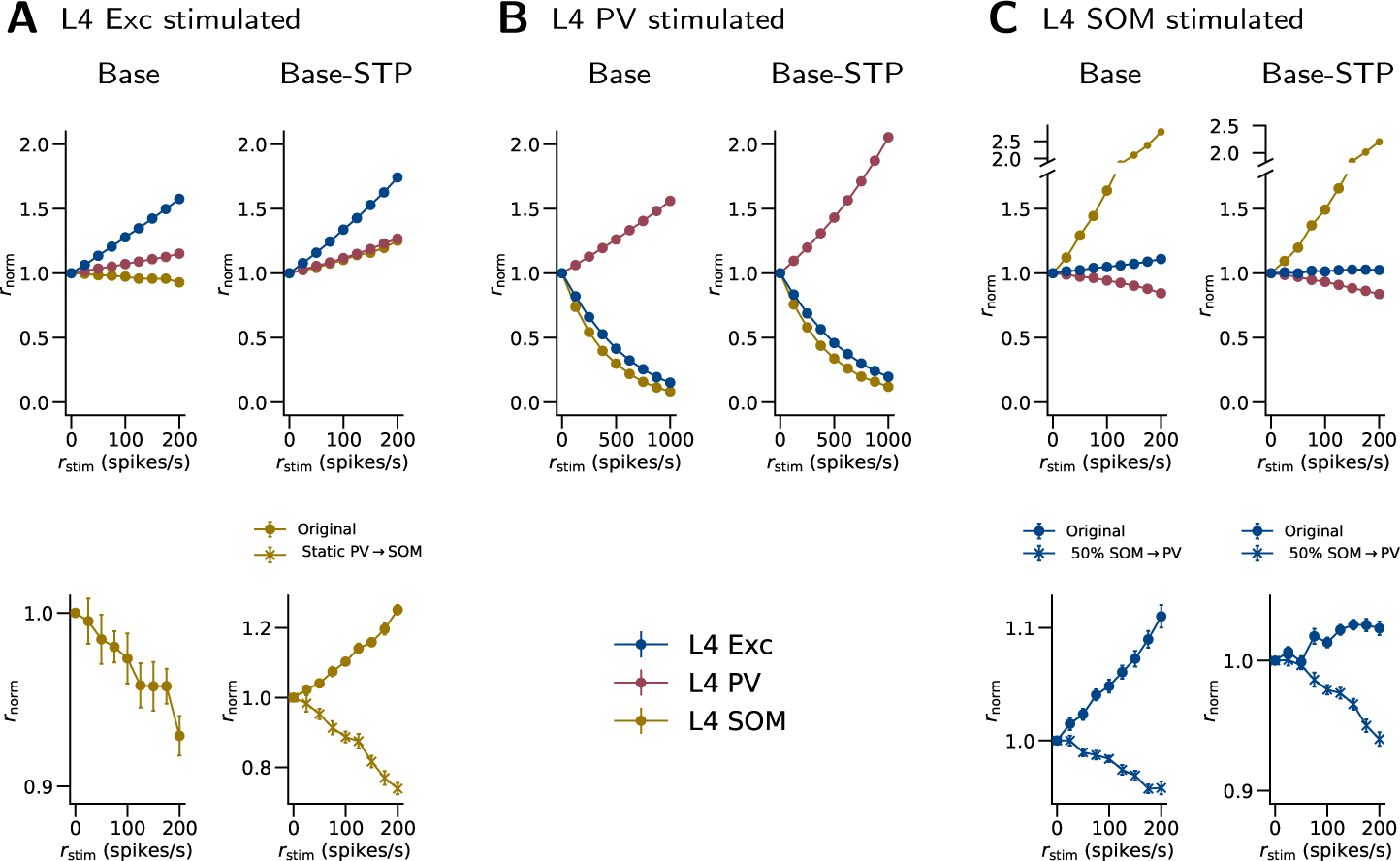
L4 network responses to cell-type-specific stimulation. Each of (A) to (C) shows the changes in population firing rates when one particular population is stimulated in L4. The notations are as in Figure 7. 50% SOM→PV: The SOM→PV connection probability is reduced to 50% of the original value.

The disinhibitory effect of VIP cells on Exc cells may require adequately active SOM cells. To examine this, we run the VIP stimulation again with a lower background input (*r*_bg_) for SOM cells. When the *r*_bg_ is lower by 200 spikes/s, the disinhibitory effect indeed disappears (Figure 7D, second row).

The contrast between inhibitory L2/3 SOM cells and disinhibitory L4 SOM cells in our model reproduces the result in the experiment by Xu et al. (2013). They suggested that this constrast is because of a difference in SOM connectivity that is also observed in their experiment, i.e., the SOM→Exc projection being stronger in L2/3 but weaker in L4 than SOM→PV. To further test this hypothesis with our model, we run the SOM stimulation again with lower L4 SOM→PV connection probability. We find that L4 SOM cells become inhibitory to Exc cells (-4.2% at *r*_stim_ = 200 spikes/s, *p <* 0.001) when the L4 SOM→PV connection probability is reduced to 50% of the original value (Figure 8C, second row). This reduced L4 SOM→PV connection probability (18.2%) is also lower than the L4 SOM→Exc connection probability (19.8%). Therefore, this result is consistent with the hypothesis proposed by Xu et al. (2013), that different SOM connectivity contributes to the observed inhibition-versus-disinhibition contrast.

In the Base-STP model, the interneurons exhibit the same inhibitory or disinhibitory effects on Exc cells (Figure 7 and 8) as their counterparts in the Base model. In L2/3, the Exc firing rate decreases significantly with PV (*−*44.6% at *r*_stim_ = 1000 spikes/s, *p <* 0.001) and SOM (*−*26.0% at *r*_stim_ = 200 spikes/s, *p <* 0.001) activation, while it increases significantly with VIP activation (+9.0% at *r*_stim_ = 200 spikes/s, *p <* 0.001). Similar to the Base model, the disinhibition of Exc cells upon VIP stimulation disappears when *r*_bg_ for SOM is lower by 400 spikes/s (Figure 7D, second row). In L4, the Exc firing rate decreases significantly with PV activation (*−*80.3% at *r*_stim_ = 1000 spikes/s, *p <* 0.001), while it increases significantly with SOM activation (+2.5% at *r*_stim_ = 200 spikes/s, *p <* 0.001). The L4 SOM cells also become inhibitory to Exc cells (*−*6.7% at *r*_stim_ = 200 spikes/s, *p <* 0.001) when the L4 SOM→PV connection probability is reduced to 50% of the original value (Figure 8C, second row).

Although mostly similar, there are still qualitative differences in the results between the two model versions. Paradoxically, the L2/3 Exc cells in the Base-STP model are initially suppressed in response to Exc stimulation (*−*8.9% at *r*_stim_ = 375 spikes/s, *p <* 0.001) but become activated with stronger stimulation, while those in the Base model are only activated (Figure 7A). We hypothesized that the STP in Exc→SOM connections contributes to this difference. Because the Exc→SOM connections are facilitated, the SOM cells may be activated more strongly upon Exc stimulation, and in turn inhibit the Exc cells (Figure 9A). To examine this hypothesis, we run the simulation with the Base-STP model but exclude the STP in Exc→SOM connections. With this change, the Exc firing rates at all stimulation levels are higher than the value at *r*_stim_ = 0 (Figure 7A, second row), supporting our hypothesis.

**Figure 9.**
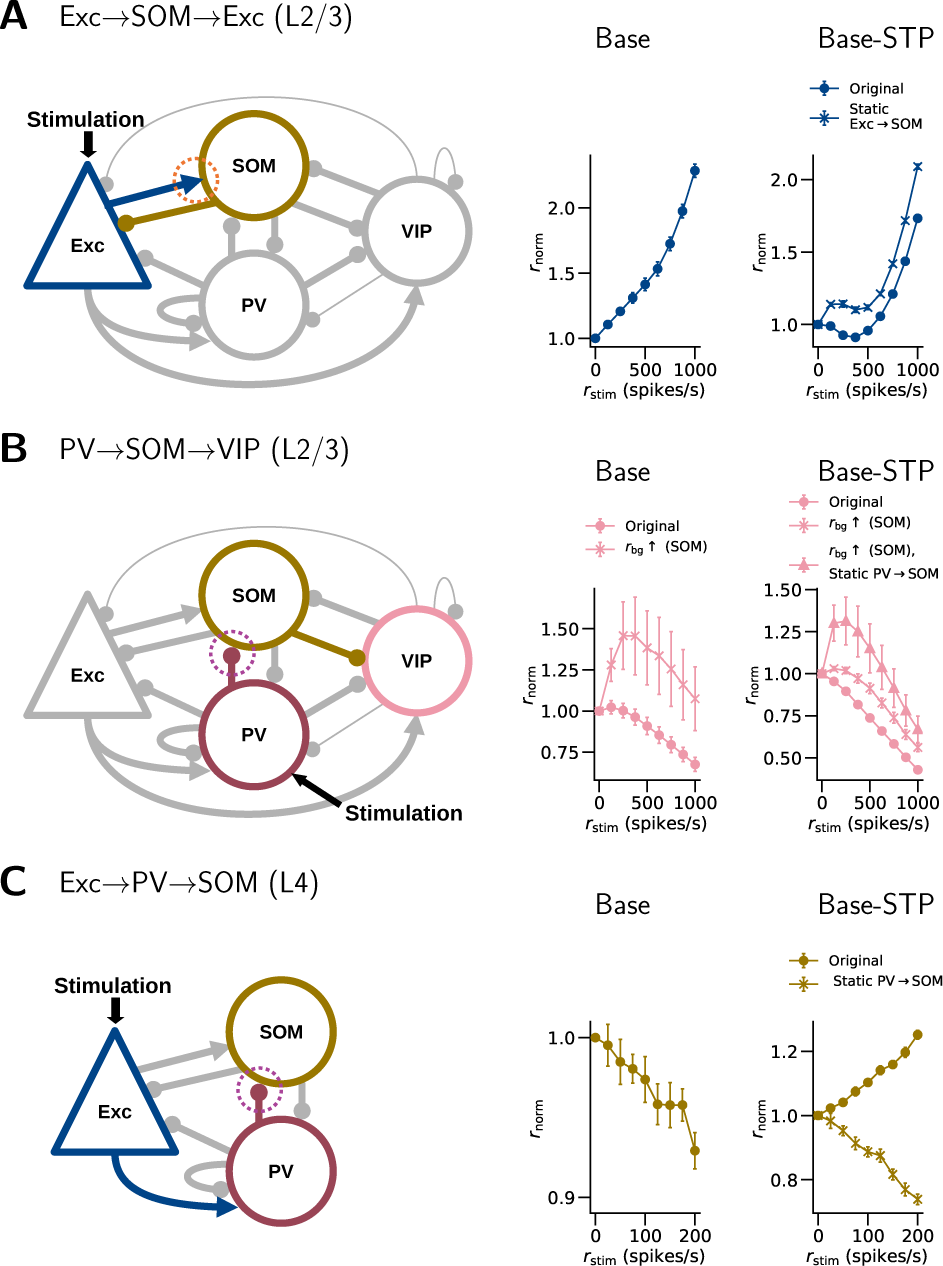
Possible pathways and STP effects underlying the differences in network responses to cell-type-specific stimulation. (A) When the L2/3 Exc cells are stimulated, a feedback inhibition of Exc cells may be mediated through the Exc→SOM→Exc pathway. Facilitation (orange dotted circle) in the Exc→SOM projection may enhance this pathway, contributing to the initial suppression of Exc cells. In support of this hypothesis, excluding the STP in the Exc→SOM projection keeps the Exc firing rate higher than the value at *r*_stim_ = 0 (Base-STP). (B) When the L2/3 PV cells are stimulated, the VIP cells may be initially disinhibited through the PV→SOM→VIP pathway, if the SOM cells are sufficiently active. Depression (purple dotted circle) in the PV→SOM projection may weaken this pathway. In support of this hypothesis, excluding the STP in the PV→SOM projection enhances the initial VIP activation (Base-STP). (C) When the L4 Exc cells are stimulated, an inhibition of SOM cells may be mediated through the Exc→PV→SOM pathway. Depression in the PV→SOM projection may weaken this pathway. In support of this hypothesis, excluding the STP in the PV→SOM projection changes the activation of SOM cells to a suppression (Base-STP). The panels show the same data as in Figure 7 and 8.

L2/3 VIP cells in the Base model show a slight increase in firing rate at the first level of PV stimulation (+2.3% at *r*_stim_ = 125 spikes/s, Figure 7B, first row). In contrast, their counterparts in the Base-STP model are suppressed at all levels of PV stimulation. We hypothesized that this initial VIP activation reflects a disinhibition of VIP cells by PV cells through a PV→SOM→VIP pathway (Figure 9B). We examine this hypothesis with modified model parameters as follows. We run the PV stimulation again with the Base model, but with a higher *r*_bg_ for SOM (+50 spikes/s) or a lower *r*_bg_ for VIP (*−*50 spikes/s). Both modifications result in a higher resting-state SOM firing rate (4.69 and 3.81 spikes/s respectively) and a significant initial VIP activation in response to PV stimulation (+45.8% and +21.1% at *r*_stim_ = 250 spikes/s, *p* = 0.038 and *p* = 0.033, respectively; The first case is shown in Figure 7B, second row). This suggests that the effect of the PV→SOM→VIP pathway is significant when SOM cells are adequately active. However, a similar modification in the Base-STP model produces a much smaller initial VIP activation (+3.0% at *r*_stim_ = 125 spikes/s, *p* = 0.02; Figure 7B, second row), even though the modification is larger (+175 spikes/s for *r*_bg_ for SOM) and produces a higher resting-state SOM firing rate (5.9 spikes/s) than in the Base model. Therefore, we further hypothesized that the STP in PV→SOM connections contributes to this difference (Figure 9B). To examine this, we run the PV stimulation with Base-STP again with higher *r*_bg_ for SOM, while excluding the STP in PV→SOM connections. In this case, the initial VIP activation (+31.3% at 250 spikes/s, *p* = 0.04; Figure 7B, second row) becomes more comparable to the modified Base model. This suggests that the excluded STP accounts for part of the observed difference.

L4 SOM cells in the Base model are suppressed in response to L4 Exc stimulation (*−*7.1% at *r*_stim_ = 200 spikes/s, *p <* 0.001; Figure 8A), while their counterparts in the Base-STP model are activated (+25.2% at *r*_stim_ = 200 spikes/s, *p <* 0.001). We hypothesized the following mechanism. In the Base model, when the L4 Exc cells are stimulated, the L4 Exc→SOM pathway is dominated by the Exc→PV→SOM pathway, which suppresses the SOM cells (Figure 9C). In the Base-STP model, however, the facilitated Exc→SOM connections reverse this dominance. To examine this hypothesis, we run the Exc stimulation with the Base-STP model again but exclude the STP in Exc→SOM connections. With this change, the SOM cells are always suppressed upon Exc stimulation (Figure 8A, second row), as in the Base model. This result supports our hypothesis.

With the same stimulation protocol, we also examined several model versions with modified parameters. We first considered a “Double-sized” model. This model version is created with twice the cell number, and the connection probability of each projection is integrated with twice the surface area. Other than these changes, the parameters remain the same. The results are well preserved in this model version, both with and without STP (Figure S5 and S6). Similarly, another version of the model with adjustments to obtain more realistic thalamocortical responses (the TC-adjusted model; see Network Responses to Thalamic Stimulation in the following) also reproduces most of the results (Figure S8 and S9). Finally, we examined the second-best-fit and third-best-fit models in the optimization process of the background input, which again yields closely similar results (Figure S12, S13, S14, and S15).

### Network Responses to Thalamic Stimulation

Figure 10 shows network responses to thalamic stimulation in terms of peristimulus time histograms (PSTH). The experimental data used for evaluation are the touch-evoked cortical responses from Yu et al. (2019). The Base model shows plausible cell-type-specific response amplitudes in L4 and L5 (Figure 10A; crosses show peaks in experimental data), and the order of their onsets and peaks also resembles the experimental data (PV→Exc→SOM; see Figure 3 in Yu et al. (2019)). However, the L2/3 responses are overestimated, and responses in all layers occur substantially earlier than in the experimental data. Starting from the Base model, we scan two sets of parameters in turn to optimize the responses.

**Figure 10.**
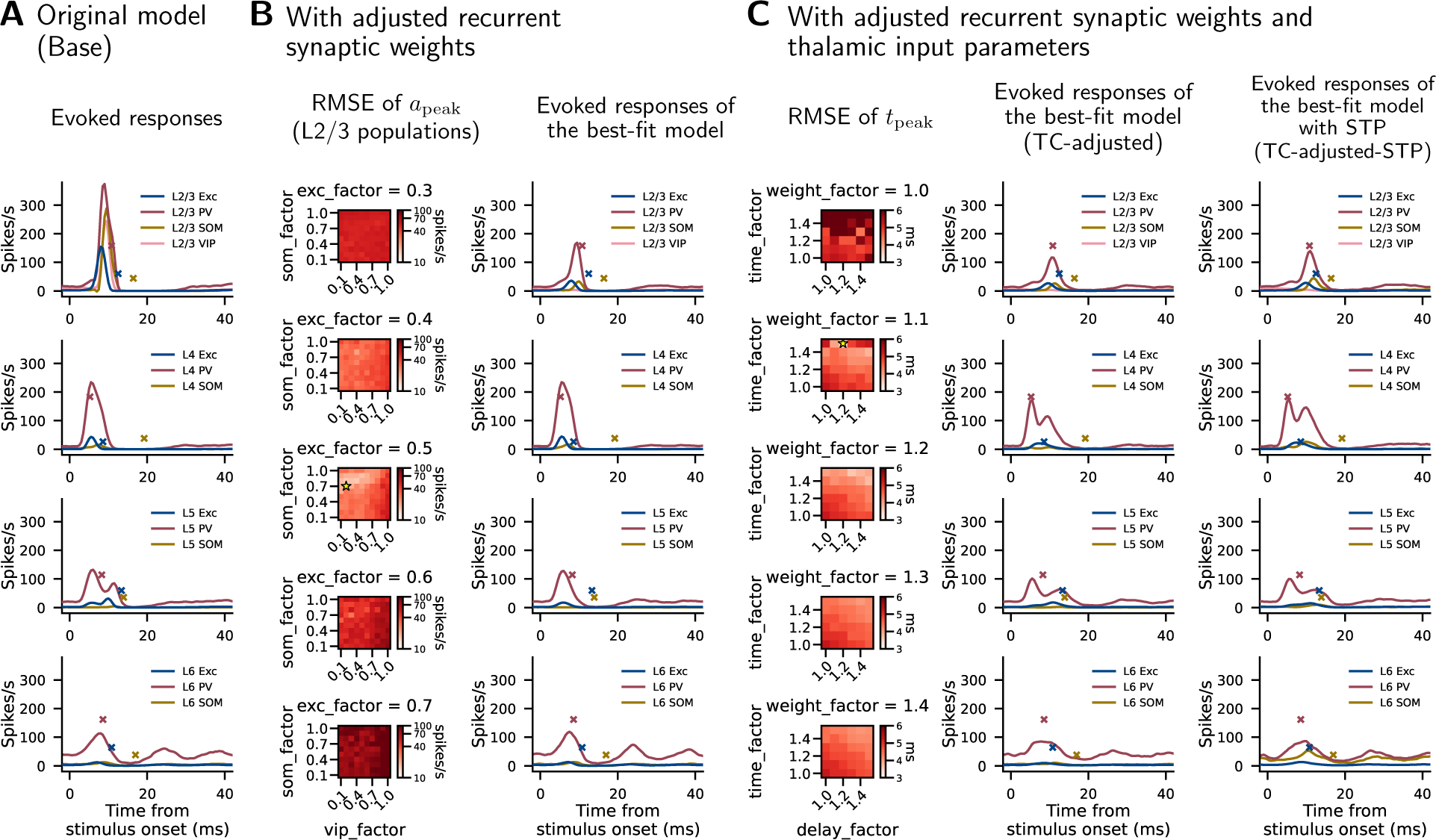
Network responses in terms of PSTH upon thalamic stimulation. (A) Responses of the original model. (B) Results with adjusted recurrent synaptic weights. Left: RMSEs of L2/3 response peak amplitudes (*a*_peak_) between simulation and experiment while scanning three factors: synaptic weights from Exc cells of all layers to L2/3 Exc (exc_factor), to L2/3 SOM (som_factor), and to L2/3 VIP (vip_factor) cells. For exc_factor, levels from 0.1 to 1.0 are scanned, but only 0.3 to 0.7 are displayed as the fitness is substantially better at these levels. Right: Evoked responses of the best-fit (with smallest RMSE) model in the scan. (C) Results with the best-fit parameters from (B) and adjusted thalamic input parameters. Left: RMSEs of response peak times (*t*_peak_) between simulation and experiment while scanning three factors: synaptic weight of thalamic input for Exc cells (weight_factor), time course of thalamic input to Exc cells (time_factor), synaptic delay of thalamic input to all cell types (delay_factor). Middle and right: Evoked responses of the best-fit model in the scan. The best-fit parameters in (B) and (C) are indicated by the stars in the heatmaps. The PSTH bin width is 1 ms. Crosses in the PSTH plots mark the peaks in the experimental data, digitized from Figure 3 in Yu et al. (2019). Data shown are the means of 10 (heatmaps) or 20 (PSTH plots) simulation instances, each with 10 repeats of stimulation.

We first explore three parameters to reduce the L2/3 responses relative to the other layers. We downscale the synaptic weights of all excitatory recurrent (intracortical) projections to L2/3 Exc, SOM, and VIP cells by factors exc factor, som factor, and vip factor respectively. PV cells are considered the main source of inhibition and therefore are not included in the parameter scan here. The heatmaps in Figure 10B show the root-mean-square errors (RMSEs) of L2/3 population-specific response peak amplitudes (*a*_peak_) between simulation and experiment while scaling these synaptic weights. The star in the heatmap represents the best-fit (i.e., with the smallest RMSE) model, and the PSTH plot shows its responses.

Next, we start from the best-fit model in Figure 10B to optimize the times of the population-specific response peak times (*t*_peak_). Because the peaks in the simulation appear to be earlier than the experimental data, we up-scale three thalamic input parameters to extend the responses as follows. (1) Time course of the thalamic input to Exc cells (time factor). This is done by extending the time course of the input (Figure 4) horizontally: let *f* (*t*) be the original time course and *f ^′^*(*t*) be the extended time course, then *f ^′^*(*t*) = *f* (*t/*time factor). (2) Synaptic weights of the thalamic input to Exc cells (weight factor). (3) Synaptic delays of the thalamic input for all cell types (delay factor). Figure 10C shows the RMSEs of *t*_peak_ between simulation and experiment while scaling these parameters (heatmaps), and the responses of the best-fit model (TC-adjusted; middle PSTH). In addition, the best-fit parameters in Figure 10C are applied to the Base-STP model to create a version with STP (TC-adjusted-STP; right PSTH). With STP, six populations show significantly different response peak amplitudes compared to the model with static synapses (TC-adjusted vs. TC-adjusted-STP, in spikes/s): L2/3 SOM (36.5 vs. 55.4, *p* = 0.007), L2/3 VIP (12.7 vs. 16.6, *p <* 0.001), L4 SOM (16.7 vs. 41.2, *p <* 0.001), L5 SOM (6.1 vs. 18.4, *p <* 0.001), L6 Exc (12.3 vs. 16.8, *p <* 0.001), and L6 SOM (20.8 vs. 77.6, *p <* 0.001).

Overall, downscaling the selected synaptic weights to appropriate levels, especially for Exc cells (exc factor), results in more plausible response amplitudes in L2/3 (Figure 10B). Then, up-scaling the three selected thalamic input parameters to appropriate levels brings the response peaks closer in time to the experimental data (Figure 10C). RMSEs of *a*_peak_ and *t*_peak_ are 117.80 spikes/s and 5.07 ms in the Base model in Figure 10A, and are improved to 36.33 spikes/s and 3.53 ms in the TC-adjusted model in Figure 10C.

The RMSE calculation for *a*_peak_ (Figure 10B) uses zero as the criterion for VIP, and the one for *t*_peak_ (Figure 10C) excludes the L2/3 VIP population. This is because there is no obvious VIP response in the experimental data (Figure 3F of Yu et al. (2019)).

Note that the parameter scan in Figure 10C includes only the thalamic input parameters; hence, the parameters of the cortical network are always the same and correspond to the best-fit model in Figure 10B. We examine the resting state and response to cell-type-specific stimulation of this cortical network, for both the static synapse (TC-adjusted) and STP (TC-adjusted-STP) versions. The results (Figure S7, S8, and S9) reproduce most of those in the original model.

## Discussion

We developed a computational model of a multi-layer cortical microcircuit incorporating three major inhibitory interneuron types, the parvalbumin-, somatostatin- and vasoactive intestinal peptide-expressing cells (PV, SOM, and VIP cells). The model is constrained by biologically plausible parameters obtained from mouse and rat somatosensory (S1) cortex on these three interneuron types. By relying exclusively on data and inferences from S1, the model’s self-consistency is enhanced. The model incorporates cell-type-specific membrane parameters, connection probabilities, and STP, which are based on experimental data. It is built with leaky integrate-and-fire neurons and is thereby computationally low-cost but still plausibly reproduces the *in vivo* resting state of different cell types. The model also reproduces known interneuron roles and provides predictions about network responses when different neuron types are stimulated as well as about the effects of cell-type-specific STP in the inhibitory control of the network. With a few adjustments, the model also shows plausible responses to thalamic input. Therefore, this model can help to theoretically and systematically study the microcircuit functions and mechanisms involving interneuron types across multiple layers of the somatosensory cortex. In this section, we discuss the links to previous studies, the limitations of the model, and potential future work.

### Model Parameters

Limited by the availability of experimental data, we use estimations and assumptions for certain model parameters. Here, we discuss the reasons for our approaches and their limitations.

Compared to the anesthetized and *in vitro* conditions, the awake state is generally considered to have lower membrane resistance (*R*_m_) and hence a shorter membrane time constant (*τ_m_*), because of frequent synaptic activity (Destexhe et al., 2003). Data indeed show that *R*_m_ of pyramidal neurons in awake mice is lower than *in vitro* (Petersen, 2017). Since *in vivo* datasets containing *τ_m_*, *R*_m_, resting and threshold potentials of all interneuron types (PV, SOM, VIP) are lacking, we used a set of *in vitro* data (Neske et al., 2015) and made an adjustment for the awake state, using experimental data on changes in *R*_m_ following transitions from Down to Up states (see Populations and Neuron Parameters, Methods). The model by Markram et al. (2015) took this issue into account by simulating with different levels of extracellular calcium concentration. Their approach requires a conductance-based neuron model with multiple ion channel types. Given that the ion channels are not explicitly modeled in this study, we compensate for the estimated Down-to-Up-state changes in *R*_m_ as an alternative approach to better approximating the *in vivo* state. As an aside, note that many *in vitro* studies use calcium concentrations very close to *in vivo* concentrations; in particular, slices showing spontaneous activity tend to have more in-vivo-like calcium concentrations than silent slices (Maksimov et al. (2018), supplementary material). However, the data from (Neske et al., 2015) we use are based on a standard calcium concentration of 2 mM for the artificial cerebrospinal fluid, which is higher than that generally measured *in vivo*. *In vivo* electrophysiological experiments with the three types of interneurons could provide the data necessary for testing and refining the corresponding assumptions.

As another adjustment to *in vivo* conditions, we have incorporated short-term plasticity (STP). While we do not find a large effect in the ongoing activity, the model reveals possible STP effects on responses to transient stimuli (see Network Responses to Cell-Type-Specific Stimulation, Results, and Network Responses to Thalamic Stimulation, Results). The similarity in resting-state activity between the models with and without STP is expected, as we adjusted the initial synaptic strengths for the STP-based model to converge to steady-state values matching the case with static synapses. Such a scheme was necessary to enable identifiability of STP-based effects separate from further differences between the two model versions there would otherwise have been. The impact of STP on transient responses is more noticeable for some populations than others. Since STP is widely present in cortical circuits, it is likely to serve a purpose even for those populations that do not show substantial effects in our model. For one, different kinds of transient stimuli beyond thalamocortical inputs are likely to meaningfully engage STP. Moreover, STP likely has meaningful effects at the level of individual synapses, neurons, and groups of neurons beyond the population level we study here. For instance, theoretical studies have demonstrated a role for STP in working memory (Romani et al., 2006; Mongillo et al., 2008; Tiddia et al., 2022) and interactions between STP and long-term plasticity (Berberian et al., 2017; Deperrois and Graupner, 2020).

Theoretically, EPSP and IPSP driving forces in neurons fluctuate with synaptic activity and associated membrane potential changes. However, *in vitro* and *in vivo* studies suggest that the effects of excitatory inputs sum close to linearly at the soma, presumably because of the isolation of individual inputs by dendritic branches and spines (Leger et al., 2005; Araya et al., 2006). Since our neuron model is a point neuron, we only model the effects of synaptic inputs on the somatic membrane potential and not local effects on the dendrites; and we therefore use current-based synapses to capture the linear summation.

For several intralaminar projections involving SOM cells, connectivity data are lacking for the primary somatosensory cortex (Figure 3). In principle, it would have been possible to fill in gaps in the data using connection probabilities from the primary visual cortex (V1), e.g., Jiang et al. (2015). However, studies have indicated significant differences in connectivity involving SOM cells between S1 and V1 (Scala et al., 2019). Therefore, we instead used averaged connection probabilities from other layers (see Probabilities of Intracortical Connections, Methods). We also assumed an equal connection probability for Exc→PV and PV→Exc projections in layer 4 and 5, where we do not find data for Exc→PV projections. This reciprocity is supported by several experimental studies (Geiger et al., 1997; Couey et al., 2013; Hu et al., 2014; Koelbl et al., 2015; Qi et al., 2017).

Because data on interlaminar connection probabilities involving specific interneuron types are lacking, we use the algorithmic estimates by Markram et al. (2015) to supplement this part of our model. Markram et al. (2015) distinguished the neuron types by morphology and provided correlations with molecular markers (Figure 2 and Table 1 in Markram et al. (2015)). As described in Methods, we calculate average connection probabilities accordingly for the projections in our model (e.g., large basket cells and nest basket cells express PV; their connection probability data were averaged and used for the PV cells in our model), although this mapping may not be very precise. With this approach, the resulting connection probabilities of interlaminar inhibitory projections are mostly *<*5 % (Figure 3). Morphological studies have suggested that, although some interneurons have axons that are mostly confined to their layers of origin, others still have significant interlaminar projections (Thomson and Bannister, 2003; Kumar and Ohana, 2008; Packer et al., 2013). Further experimental data on functional connectivity can verify if the connection probabilities in our model fairly represent these interlaminar projections and improve the estimates where necessary.

It should further be noted that connection probabilities obtained from paired recordings in brain slices, such as those included in this study, may suffer from underestimation because of truncation of axons and dendrites during slice preparation (Lefort et al., 2009; Stepanyants et al., 2009; van Albada et al., 2022). In this regard, more experimental data combining advanced neuron classification and connectivity reconstruction methods (Kebschull et al., 2016; Gouwens et al., 2020) may improve the accuracy of the connectivity in our model.

Fast-spiking (FS) cells are taken as PV cells to obtain data on connection probability or STP in some cases, where molecular marking is not done in the experiments (Kapfer et al., 2007; Hu et al., 2011; Ma et al., 2012). The close relation between FS cells and PV cells has been well established in previous rodent studies. This includes experiments for several cortical areas such as somatosensory cortex, visual cortex, and frontal cortex, by means of electrophysiological recordings and antibody labeling of individual neurons (Kawaguchi and Kondo, 2002; Chattopadhyaya et al., 2004; Koelbl et al., 2015). Therefore, we consider the mapping of FS data to PV cells to be reliable.

For simplicity, we do not take into account differences in single-neuron firing patterns between the cell types. While these probably affect network dynamics to some extent, the effects of single-neuron firing patterns may be limited on the network level depending on the dynamical state. For instance, when the network is squarely in the asynchronous regime, the effects of single-neuron bursting may not be obvious (Sahasranamam et al., 2016). The investigation of the effects of diverse single-neuron firing patterns in the present model is left to future work.

The firing rates of the background inputs (*r*_bg_) are optimized to obtain plausible resting-state population firing rates. We found that the resulting *r*_bg_ shows a pattern of PV *>* Exc *>* VIP *>* SOM, which is also found in some experimental data of long-range inputs to these cell types in S1 in recent studies (Martinetti et al., 2022; Naskar et al., 2021). This indicates that our optimization is biologically plausible at least in this respect. The level of background input is also similar to that in the model of Potjans and Diesmann (2014). The effective strength of the external drive onto excitatory cells is 5000 spikes/s *×* 0.5 mV/spike = 2500 mV/s, similar to that in Potjans and Diesmann (2014), which is 2000 *×* 8 spikes/s *×* 0.15 mV/spike = 2400 mV/s. This similarity also holds for the average onto the inhibitory cells (2091 vs. 2220 mV/s). Compared to Potjans and Diesmann (2014), we use a stronger unitary weight, which is based on data from paired recording experiments, but the summed strength of the external input is approximately conserved. Note, however, that the spatial extent of our model is 0.06 mm^2^ as opposed to the 1 mm^2^ of the Potjans and Diesmann (2014) model, so that our circuit includes a smaller percentage of the sending neurons, and a larger percentage of the input to the neurons is provided by the external drive. That the external drive is nevertheless close to that in their model appears mostly related to the different neuron parameters, which are cell-type-specific in our model but were taken to be cell-type-independent in the model of Potjans and Diesmann (2014).

Incorporating further interneuron diversity should be considered as a next step. In addition to connectivity, SOM and PV cells also have very diverse morphologies across layers (Muñoz et al., 2017; Feldmeyer et al., 2018; Gouwens et al., 2019). Although the vast majority of VIP cells are located in L2/3, smaller numbers still exist in the deeper layers which show different dendritic and axonal projection patterns (Prönneke et al., 2015). It was also revealed recently that a possible subgroup of PV cells can mediate a thalamus-driven disinhibition in L4 (Hua et al., 2022). How this diversity of interneurons contributes to inhibitory control and computation in the cortical column can be investigated in future by extending and refining our model.

### Short-Term Synaptic Plasticity

By fitting post-synaptic potentials to the model of Tsodyks et al. (2000), we systematically determined cell-type-specific parameters of short-term plasticity (STP) that may be useful for future modeling studies. As explained in Methods, the synaptic weights of the model with STP (Base-STP) are scaled so that its resting-state synaptic weights are close to those set for the model with static synapses (Base). This ensures that the two model versions have similar resting states so that responses to stimulation can be fairly compared. This also means that the Base-STP model reproduces the *in vivo* state just as well as the Base model, but under synaptic depression and facilitation. This allows computational studies of STP effects. Since the *in vivo* synaptic weights before depression and facilitation are difficult to measure or estimate, incorporating STP parameters as we have done provides a tool complementing what is possible experimentally.

Similar to the Base model, the asynchronous irregular (AI) activity in the Base-STP model is consistent with that observed *in vivo*, as assessed using criteria compiled by Maksimov et al. (2018). The main difference is that the average L6 pairwise spike count correlation in the Base-STP model is more than twice as large as that in the Base model (Figure 6B). Although synchronized oscillatory activity in the cortex has sometimes been considered to be generated by intrinsic cell membrane mechanisms (Silva et al., 1991; Le Bon-Jego and Yuste, 2007; Hayut et al., 2011), it has also been linked to the specific connectivity and synaptic dynamics of PV and SOM cells (Le Bon-Jego and Yuste, 2007; Fanselow et al., 2008; Chen et al., 2017; Funk et al., 2017; Veit et al., 2017; Domhof and Tiesinga, 2021). Future work can investigate whether synaptic dynamics indeed increases L6 correlations in *in vivo* circuits as predicted by our model.

### Cell-Type-Specific Stimulation

Results of cell-type-specific stimulation in L2/3 shows that PV and SOM cells are inhibitory and VIP cells are disinhibitory to the Exc cells (Figure 7), consistent with expectations based on the experimental literature (Beierlein et al., 2003; Kapfer et al., 2007; Silberberg and Markram, 2007; Lee et al., 2013; Pfeffer et al., 2013; Pi et al., 2013; Hu et al., 2014; Karnani et al., 2014, 2016a; Naka and Adesnik, 2016; Yavorska and Wehr, 2016). Furthermore, several model versions with modified parameters also reproduce these results (Figure S5, S6, S5, S6, S12, S13, S14, S15), confirming the robustness of the model.

Several results in L2/3 may be of interest for further study: (1) With Exc cell stimulation, the VIP cells are suppressed in both the Base and Base-STP models. This may be due to the activation of SOM cells and a consequent SOM→VIP inhibition, as this SOM→VIP projection is supported by experimental data (Karnani et al., 2016b) and implemented in our model as well. (2) With Exc cell stimulation, the Exc cells in the Base-STP model are paradoxically suppressed initially with weak stimulation strengths, then become activated again with stronger stimuli. The initial dampening may be due to the facilitated Exc→SOM projection, which could enhance the activation of SOM cells and in turn suppress the Exc cells (Figure 9A). (3) VIP cells in the Base model tend to be initially activated in response to PV stimulation. We hypothesized that this is because the direct PV→VIP inhibition is weaker than the disinhibition of VIP cells through a PV→SOM→VIP pathway. The depressing PV→SOM projection in the Base-STP model may reduce this dominance, and hence weaken the disinhibition of VIP cells (Figure 9B). We tested the hypothesized STP effects in (2) and (3) by excluding the STP of corresponding projections and obtained supporting results (Figure 7A and B). However, there is still a possibility that other factors also contribute in parallel to the observed differences, e.g., STP of other projections. These predictions should be examined in further experimental or theoretical studies.

With L4 cell-type-specific stimulation in our model, we observe the following: (1) L4 SOM cells show a disinhibitory effect on L4 Exc cells, in contrast to L2/3 SOM cells, which show an inhibitory effect (Figure 8). We believe this reflects the higher SOM→PV connection probability in L4 than in L2/3 in our model (36.30% vs. 11.81%; see Figure 3), consistent with the experiment by Xu et al. (2013) (see Comparisons with Relevant Models in the following). The result shows that our model can reflect layer-specific roles of SOM cells. (2) L4 Exc stimulation suppresses L4 SOM cells in the Base model, but activates them in the Base-STP model. The suppression in the Base model may be because the Exc→SOM projection is dominated by the Exc→PV→SOM pathway, while the depressing PV→SOM projection in the Base-STP model may override this dominance (Figure 9C). We tested this hypothesis by excluding the STP of the PV→SOM projection and obtained supporting results (Figure 8A). Like those in L2/3, these results are worth further experimental or theoretical studies.

In both model versions, the sensitivity of Exc, PV, SOM, and VIP cell activity to the stimulation of their own populations is very different. The slopes of the normalized stimulation-induced firing rates in L2/3 are ordered as SOM*>*VIP*>*Exc*>*PV (Figure 7 and 8). To the best of our knowledge, the available experimental data do not allow for direct comparison with this result, for lack of analogous cell-type-specific stimulation experiments. Therefore, the result provides another prediction, which can be examined by future experiments distinguishing the given cell types. A possible cause for the observed difference in sensitivity is the cell-type specificity of the membrane time constants in L2/3, which also follow the order SOM*>*VIP*>*Exc*>*PV (Table 2). This is because a larger membrane time constant increases the area under the PSPs onto the cell and hence the probability of the cell being brought to fire. With further analytical methods, our model can help to predict other neuronal or circuit-level factors behind this result.

### Thalamic Stimulation

We assess the capability of our model to simulate sensory responses by comparing with the *in vivo* data (Yu et al., 2019). In response to thalamic stimulation, the Base model shows a few plausible cell-type-specific responses but still has substantial discrepancies (Figure 10A). There may be a few causes for these discrepancies.

To determine the cause for the overestimated L2/3 responses, we tested the thalamic stimulation in a model version without L4 Exc→L2/3 Exc connections and found that the L2/3 responses became much smaller than the *in vivo* data (data not shown). This suggests that the feedforward excitatory projections from L4 to L2/3 are the main source of excitation for the L2/3 responses, transmitting the thalamic input indirectly. As described in Methods, we based the connection probabilities and STP of Exc→Exc projections on layer-specific experimental data. However, the STP of all other projections and all synaptic weights in our model are not layer-specific. The impact of layer specificity of these parameters may be investigated as corresponding data becomes available. To improve the L2/3 responses in the present study, we perform a parameter scan including synaptic weights of recurrent excitatory projections to the L2/3 populations. The best-fit model in this scan shows plausible L2/3 response amplitudes (Figure 10B).

Also the timings of thalamocortical responses are not perfectly predicted. In part, this may reflect the fact that some external inputs associated with sensory responses are missing in our model. For example, higher-order or non-specific thalamic nuclei such as the posterior medial nucleus (POm) may contribute to longer responses (Zhang and Bruno, 2019). Indirect or feedback inputs from other barrel columns or even cortical areas may also substantially extend the responses (Aronoff et al., 2010; Minamisawa et al., 2018). These inputs have not been considered here as we only incorporate the ventral posteromedial nucleus (VPM). In the absence of the corresponding parameter values, we examine the possibility of extending the responses by scanning three thalamic input factors. Figure 10C shows that up-scaling the three selected factors helps to bring the response timings closer to the experimental data. This result suggests that incorporating further external inputs can help to similarly reproduce the experimentally observed response properties.

We also compare the TC-adjusted model with and without STP (Figure 10C). The response peak amplitudes are different in a few populations, mainly the SOM cells. This may reflect the higher resting-state SOM firing rates in the Base-STP model (Table 5). On the other hand, almost all Exc and PV populations, comprising most of the neurons, do not show a significant difference. As in the case of cell-type-specific stimulation, we believe the overall similarity is associated with similar resting states. However, it may also be related to the transient nature of the stimulation, which diminishes the STP effects that need a longer time to manifest. Given the association between STP and frequency responses of neurons (Beierlein et al., 2003), whether other qualitative differences will appear with different types of stimulation (e.g., longer duration or higher frequency) can be the next subject of study.

### Comparisons with Relevant Models

In recent years, increasing attention has been devoted to incorporating the major interneuron types in modeling studies to understand cortical microcircuit dynamics and signal processing (Litwin-Kumar et al., 2016; Yang et al., 2016; Del Molino et al., 2017; Lee and Mihalas, 2017; Lee et al., 2017; Hertäg and Sprekeler, 2019; Mahrach et al., 2020; Sanzeni et al., 2020; Borges et al., 2022; Hertäg and Clopath, 2022; Guo and Kumar, 2023; Moreni et al., 2023; Wagatsuma et al., 2023). In particular, several multi-layer models of cortical areas S1 and V1 incorporating multiple interneuron types have been developed (Markram et al., 2015; Billeh et al., 2020; Borges et al., 2022; Moreni et al., 2023). Morphological or physiological data from S1 (Markram et al., 2015; Billeh et al., 2020; Borges et al., 2022) or V1 (Billeh et al., 2020; Moreni et al., 2023) were used to derive neuron and connectivity parameters and to establish models with Leaky-Integrate-and-Fire (LIF) (Markram et al., 2015; Billeh et al., 2020; Moreni et al., 2023) or multi-compartment (Markram et al., 2015; Billeh et al., 2020; Borges et al., 2022) neurons. With these models, network synchrony (Markram et al., 2015), oscillatory activity (Moreni et al., 2023), and selective sensory responses (Billeh et al., 2020) were studied. Moreni et al. (2023) showed how gamma oscillations around 30 Hz may arise in the presence of synaptic plasticity. Our model displays higher-frequency gamma oscillations that have not been observed experimentally in this form. The frequency and amplitude of gamma oscillations in models based on balanced random networks depends on a multitude of factors, the examination of which is beyond the scope of this study. For preliminary work on a thorough investigation of this issue, see Essink et al. (2020).

Our model is adapted from Potjans and Diesmann (2014) with major parameter changes. As described, we base the model exclusively on data from mouse and rat somatosensory cortex and incorporate three interneuron types and their cell-type-specific STP. The model of Potjans and Diesmann (2014), which groups the inhibitory interneurons into a single population per layer, already reproduces several aspects of resting-state activity and sensory responses. Thus, distinguishing the interneuron types is not necessary to account for major properties of the low-rate asynchronous irregular resting state. However, distinguishing these cell types enables relationships between the structure and dynamics of the cortical microcircuitry to be explored in more detail.

Based on experimental data on rat somatosensory cortex, Markram et al. (2015) constructed an *in silico* cortical microcircuit with a multicompartmental and conductance-based neuron model. As mentioned, they estimated microcircuit activities under different extracellular calcium concentrations to mimic differences between *in vitro* and *in vivo* data. The authors also simulated a thalamic activation of the microcircuit and reproduced the response pattern of cortical neurons in experimental data (Figure 17 in Markram et al. (2015)). Compared to the highly detailed model of Markram et al. (2015), our model can be used to more easily simulate and mechanistically analyze cell-type-specific network dynamics, with smaller computational resources.

Mahrach et al. (2020) studied the paradoxical effect, where stimulation reduces rather than increases the firing rate of inhibitory cells (Tsodyks et al., 1997; Murphy and Miller, 2009; Ozeki et al., 2009). They stimulated PV cells *in vivo* in mouse anterior lateral motor cortex and barrel cortex and compared the results with those of a computational model. Their model is able to reproduce the paradoxical effect found in experimental data and provides predictions on the underlying parameter values. Specifically, for their “Model 1”, the paradoxical effect of PV cells depends on *J*_SV_(*J*_EE_*J*_VS_ *− J*_ES_*J*_VE_), where *J*_XY_ stands for the interaction strength from population Y to X (X, Y *∈ {*E, S, V*}*; E, S, V stand for Exc, SOM, and VIP cells, respectively). In their theory, if *J*_EE_ is small enough to make *J*_SV_(*J*_EE_*J*_VS_ *− J*_ES_*J*_VE_) negative, the PV cells should show the paradoxical effect. Here, we compare the L2/3 part of our model with their Model 1, which both have Exc, PV, SOM, and VIP cells. With our original parameters, the paradoxical effect is absent (Figure 7B and S16A). As an effort to eliminate differences between their model and ours that could block the paradoxical effect, we tested our L2/3 network (1) with *J*_EE_ being zero or very weak (down to 1/128 of the original), which predicts a paradoxical effect in their model, (2) without the extra projections (VIP→Exc, VIP→PV, VIP→VIP) and layers (L4 to L6) that are not present in their model, (3) with very weak stimulation strength for PV cells (down to 1 spike/s with a PSP amplitude of 0.5 mV), and (4) with a double-sized model. We did not observe a paradoxical effect for (1) to (3) but found a slight initial decrease in PV activity (-1.2% and -1% at stimulation strengths of 12.5 and 25 spikes/s, respectively) with the double-sized model (Figure S16B). Both Mahrach et al. (2020) and another similar study by Sanzeni et al. (2020) indicated that a network size smaller than the ones they studied (76,800 neurons and a cortical surface area of 500+ *µ*m diameter, respectively) may fail to show a paradoxical effect. The fact that we observe a slight paradoxical effect in an up-scaled model is consistent with their inference, although further studies may be required to reveal the mechanisms in detail.

Bos et al. (2020) used a spiking neuron model to analyze the influence of PV and SOM cells and showed that the role of SOM cells depends on two particular pathways: When the SOM→PV→Exc pathway dominates, SOM cells are disinhibitory, whereas when SOM→Exc and PV→PV dominate, SOM cells are inhibitory. Experimental results indicate that SOM cells can indeed be inhibitory or disinhibitory depending on the circuitry, as they show inhibitory effects in L2/3 but disinhibitory effects in L4 (Xu et al., 2013). We tested the pathway dependence of these effects in our model, considering the L4 SOM cells. With our original connectivity, the L2/3 SOM cells are inhibitory and L4 SOM cells are disinhibitory (Figure 7 and 8), which is consistent with Xu et al. (2013). By changing the L4 SOM→L4 PV connection probability, we also found that the L4 SOM cells can be either inhibitory or disinhibitory (Figure 8), reproducing the findings of Bos et al. (2020). These results are consistent with the theories and experimental observations that layer- and population-specific connectivity contributes to the layer-specific roles of SOM cells (Xu et al., 2013; Muñoz et al., 2017; Bos et al., 2020).

It should be noted that the layer-specific roles may ultimately depend on the neuronal subgroups involved, especially Martinotti vs. non-Martinotti SOM cells (Xu et al., 2013; Muñoz et al., 2017; Emmenegger et al., 2018). Such SOM cell subgroups are not explicitly considered in our model yet. Nevertheless, the layer-specific connectivity data we used are associated with these subgroups and can approximately reflect their laminar distributions. Our references include L2/3 Martinotti cells (Xu et al., 2013; Walker et al., 2016), L4 Non-Martinotti cells (Xu et al., 2013; Scala et al., 2019), and L5 Martinotti and Non-Martinotti cells (Hilscher et al., 2017; Nigro et al., 2018). An adaptation of our model that explicitly incorporates SOM cell subgroups will allow further studies on this topic. Further refinements may include additional interneuron (sub)types, such as those in layer 1 (Egger et al., 2015), which is not yet considered in our model.

### Outlook

Our model can be used as a convenient template for computational studies of complex mechanisms such as the effects of neuromodulators on sensory signal processing. Being based on the LIF neuron model, mean-field analyses can provide mechanistic explanations of the population-level dynamics, and predict network activities with different parameters or inputs without running simulations (Layer et al., 2022). This can help to facilitate the exploration of the effects of cell and connection properties and examine the hypotheses we have proposed on the main pathways underlying the suppression and enhancement of cell-type-specific activity (see Cell-Type-Specific Stimulation in the preceding). In particular, a mean-field analysis incorporating synaptic STP (Romani et al., 2006) may help reveal the mechanisms underlying the association between STP and the roles of different types of interneurons. A further future step can be to use data from public anatomical databases on barrel cortex, such as BarrelCortexInSilico (Udvary et al., 2022)^5^, that can help improve the model with more details on connectivity and other parameters. These further developments will allow more refined and systematic explorations of the roles played by different types of interneurons in cortical circuits.

## Author contributions statement

H.J. and S.v.A. conceived the study. H.J., G.Q. and D.F. contributed to the interpretation of the experimental data. H.J. created and implemented the model, performed the simulations, produced the figures, and wrote the first manuscript draft. H.J. analyzed and refined the model under supervision of S.v.A. All authors contributed to the final manuscript.

## Acknowledgments

This project received funding from the European Union’s Horizon 2020 Framework Programme for Research and Innovation under Specific Grant Agreement No. 945539 (Human Brain Project SGA3) and the European Union’s Horizon Europe Programme under the Specific Grant Agreement No. 101147319 (EBRAINS 2.0 Project).

We thank Dr. Alexander van Meegen for help with the implementation of the connection probability integration and Dr. Jorge F. Mejias for discussions on the fitting of the synaptic weights for the model with STP. We further thank Jasper Albers and Anno Kurth for determining decay constants by fits to data from Perin et al. (2011) and Packer and Yuste (2011).

## Data and code availability

The model parameters and evaluation criteria described in this article are derived from publicly available research literature and databases. The code used for data acquisition and adaptation and model implementation are available at https://zenodo.org/doi/10.5281/zenodo.10069507.

## Footnotes

1 https://nest-simulator.readthedocs.io/en/latest/models/iaf_psc_exp.html

1 Exponential decay constants.

2 Latencies from the presynaptic spike to the start of postsynaptic current.

2 https://bbp.epfl.ch/nmc-portal/welcome.html https://bbp.epfl.ch/nmc-portal/assets/documents/static/Download/layer_download.json https://bbp.epfl.ch/nmc-portal/assets/documents/static/Download/pathways_anatomy_factsheets_simplified.json

3 https://www.nest-simulator.org/

4 https://www.openmp.org/

5 https://cortexinsilico.zib.de/

