## Supplementary Material for "A Layered Microcircuit Model of Somatosensory Cortex with Three Interneuron Types and Cell-Type-Specific Short-Term Plasticity"

### Relative Quantities of Interneuron Types

The relative quantities of PV, SOM, and VIP cells ($f_{\mathrm{PV}}, f_{\mathrm{SOM}}, f_{\mathrm{VIP}}$) of each layer are obtained from Lee et al. (2010) with WebPlotDigitizer[1] as follows. The bar lengths for PV, SOM, and 5-$\mathrm{HT_{3A}}$R in Figure 2D were measured, which represent their relative quantities ($f_{\mathrm{PV}}, f_{\mathrm{SOM}}, f_{\mathrm{5HT_{3A}R}}$). Here, the relative quantity of PV/SOM (positive for both markers) is distributed equally to PV and SOM. Then, the bar length for VIP in Figure 4D was measured, which represents the fraction of VIP in 5-$\mathrm{HT_{3A}}$R. $f_{\mathrm{5HT_{3A}R}}$ is multiplied by this VIP fraction to obtain the relative quantity of VIP ($f_{\mathrm{VIP}}$).

### Postsynaptic Currents

Figure S1 shows the postsynaptic currents (PSCs) as the synaptic weights of the model. Figure S1A shows the PSCs of the model with static synapses (Base), which are calculated with Equation (2) using the cell-type-specific membrane parameters and defined PSPs (Table 2 and 3 in the main article). For each projection, the PSCs follow a log-normal distribution across connections, and the mean and standard deviation have the same magnitude.

For the model with STP (Base-STP), the PSCs are dynamic. As in the Base model, the initial PSCs are set according to log-normal distributions. However, the PSCs of different intracortical projections are scaled individually, such that the resting-state PSCs (Figure S1B) approximate those of the Base model (Figure S1A). For the thalamic input ($\mathrm{Exc_{th}}$), the initial PSCs in the Base-STP model are taken to be the same as the corresponding weights in the Base model (Figure S1A, the column for $\mathrm{Exc_{th}}$).

[1] https://automeris.io/WebPlotDigitizer/

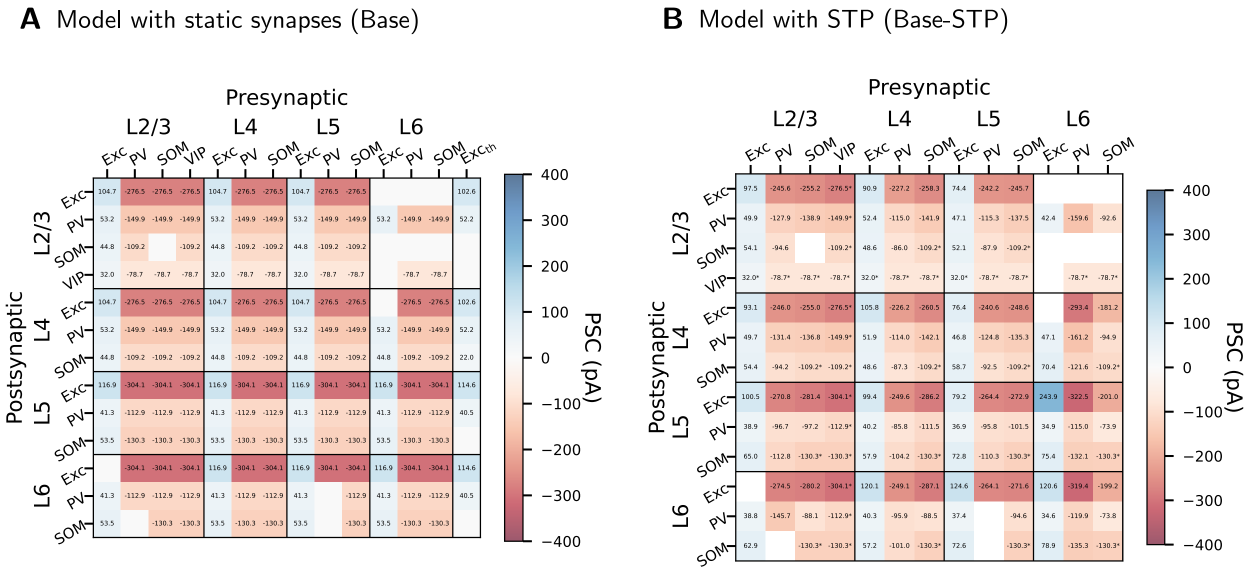


**Figure S1: Postsynaptic current in pA.** (A) Model with static synapses (Base). For each projection, the PSCs are log-normally distributed across connections. Values shown are the means, and the standard deviation for each projection has the same magnitude as the mean. These PSCs are determined by the membrane parameters (Table 2 in the main article) and PSPs (Table 3 in the main article) defined for the model. Blanks: The connection probability is zero. (B) Model with STP (Base-STP). PSCs in the Base-STP model are dynamic. Values shown are the means across 20 simulation instances of the data recorded from $t = 10\,\mathrm{s}$ to $t = 11\,\mathrm{s}$. Asterisks: Projections in the Base-STP model that remain static and use the same PSCs as their counterparts in (A).

### Derivation of Connection Probabilities

For each projection, we first estimate a theoretical zero-distance connection probability ($p_0$) from the experimentally observed probability ($P_{\text{exp}}$). We assume that connection probabilities decay exponentially so that the average connection probability within a hypothetical sphere (where recordings took place) can be expressed in terms of $p_0$ and other variables with Equation (S1) (adapted from Maksimov et al. (2018)). In other words, the unknown underlying $p_0$ of an experimentally observed average connection probability $P_{\text{exp}}$ can be estimated with the reversed form, Equation (S2):

$$P_{\text{exp}} = \frac{9}{\pi}\frac{p_0}{\text{R}^6}\int_0^{\text{R}}\int_0^{\text{R}}\int_0^{\pi} e^{\frac{-\sqrt{r_i^2+r_j^2-2cos(\alpha)r_ir_j}}{\lambda}} r_i^2 r_j^2 \text{d}r_i \text{d}r_j \text{d}\alpha \tag{S1}$$

$$p_0 = \frac{\pi}{9}\frac{P_{\text{exp}}\text{R}^6}{\int_0^{\text{R}}\int_0^{\text{R}}\int_0^{\pi} e^{\frac{-\sqrt{r_i^2+r_j^2-2cos(\alpha)r_ir_j}}{\lambda}} r_i^2 r_j^2 \text{d}r_i \text{d}r_j \text{d}\alpha} \tag{S2}$$

where $r_i$ and $r_j$ represent the distances of neurons i and j from the center of the hypothetical sphere, and $\alpha$ represents the angle formed between the two neurons and the center. R is a constant representing the radius of the sphere, which is determined by the experiment: half the maximum inter-soma distance of recordings in each experiment. $\lambda$ is the exponential decay constant of connection probability, and we use 208.3 and 123.9 $\mu$m for excitatory and inhibitory connections respectively (Figure S2), according to numerical fits of experimental data (Perin et al., 2011; Packer and Yuste, 2011) by Jasper Albers and Anno Kurth (personal communication). If multiple experiments are available for estimating $p_0$ for a projection, the average $p_0$ is used for this projection.

Second, we define the model dimensions as layers of cylinders (Figure 2 in the main article) with a radius of 138.2 $\mu$m, to fit a 200×300 $\mu$m area suggested by Petersen (2019), and layer thicknesses according to Lefort et al. (2009). $p_0$ and Equation (S3) can then be used to derive the connection probability ($P$) that fits these dimensions:

$$P = \frac{\int\int p_0 e^{\frac{\sqrt{-\rho_i^2+\rho_j^2-2\rho_i\rho_j\cos(\phi_i-\phi_j)+(z_j-z_i)^2}}{\lambda}} \text{d}v_i \text{d}v_j}{\int\int \text{d}v_i \text{d}v_j} \tag{S3}$$

where $\rho_i$ and $\rho_j$ represent the radial distances, $\phi_i$ and $\phi_j$ the radial angles, and $z_i$ and $z_j$ the vertical positions of neuron i and j in the cylindrical coordinate system, and the constant $\lambda$ is the same as in Equation (S1). Equation (S4) is used for the integration over volumes ($v_i$, $v_j$) in equation (S3):

$$\int\int \text{d}v_i \text{d}v_j = \int_0^{\text{R}} \rho_i \text{d}\rho_i \int_0^{\text{R}} \rho_j \text{d}\rho_j \int_0^{2\pi} \text{d}\phi_i \int_0^{2\pi} \text{d}\phi_j \int_{\text{Z}_1}^{\text{Z}_2} \text{d}z_i \int_{\text{Z}_3}^{\text{Z}_4} \text{d}z_j \tag{S4}$$

where $\text{R} = 138.2$ $\mu$m, and $\text{Z}_1, \text{Z}_2, \text{Z}_3, \text{Z}_4$ are the vertical positions of the lower and upper boundaries of the corresponding layers (Figure 2 in the main article). With these, the $P$ that fits the dimensions is calculated. Neurons are then connected via Bernoulli trials with probability $P$. The resultant $P$ of all projections are listed in Figure 3 in the main article.

For the interlaminar connections, there are two sets of data (Lefort et al., 2009; Markram et al., 2015), for which the maximum horizontal sampling distance is available. Because of the vertical distance between the layers, we assume that the sampling space for these data is cylindrical, and use Equation (S3) and (S4) to obtain $p_0$ and $P$. The approach is the same as described above, but because the assumed space is cylindrical, $p_0$ is estimated with a reversed form of Equations (S3) and (S4) (with R=the maximum horizontal distance/2), instead of (S2).

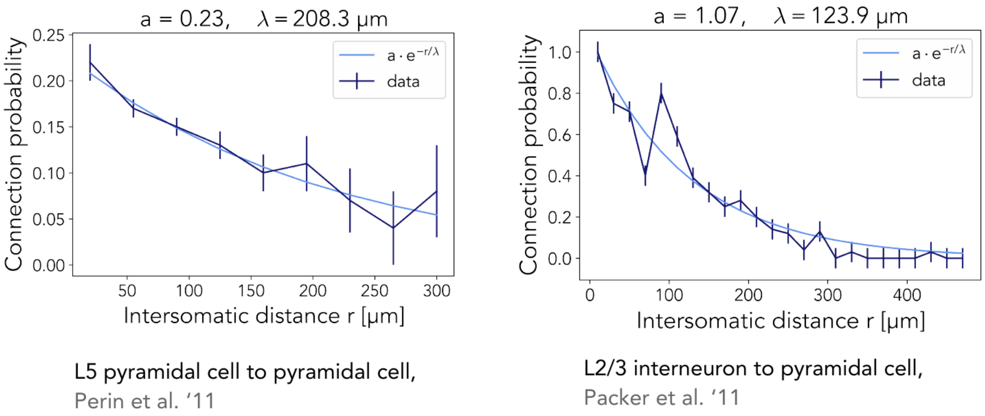


**Figure S2: Fitting of exponential decay constants of excitatory and inhibitory connection probabilities.** Data fitted by Jasper Albers and Anno Kurth, with the experimental data from Perin et al. (2011) and Packer and Yuste (2011).

### Optimization of the Background Input

The firing rates ($r_{\mathrm{bg}}$) of the background inputs for Exc, PV, SOM, and VIP cells are optimized to obtain realistic population firing rates in the model. The fitness is quantified by the root-mean-square error (RMSE) of L2/3 and L4 relative population firing rates between simulated and experimental data:

$$\mathrm{RMSE} = \sqrt{\frac{\sum_{x=1}^{\mathrm{n_{pop}}} \left(\frac{r_{\mathrm{sim},x} - r_{\mathrm{exp},x}}{r_{\mathrm{exp},x}}\right)^2}{\mathrm{n_{pop}}}}, \quad r_{\mathrm{sim},x} = \frac{\sum_{y=1}^{\mathrm{n_{sim}}} r_{\mathrm{sim},x,y}}{\mathrm{n_{sim}}},$$

where $r_{\mathrm{sim},x,y}$ is the firing rate of population $x$ in simulation $y$, $r_{\mathrm{sim},x}$ is the mean firing rate of population $x$ across all simulations, $r_{\mathrm{exp},x}$ is the firing rate of population $x$ from the experimental data, $\mathrm{n_{pop}}$ is the number of populations, and $\mathrm{n_{sim}}$ is the number of simulations performed.

The parameter scan for this optimization is performed separately for the models with static synapses and with STP. Figure S3 shows examples of the scans performed. The best-fit $r_{\mathrm{bg}}$ for static synapses are 5000 (Exc), 6900 (PV), 2500 (SOM), and 3300 (VIP) spikes/s. Those for STP are 5000 (Exc), 6800 (PV), 2400 (SOM), and 3600 (VIP) spikes/s. These values are used in a layer-independent manner for the Base and Base-STP models respectively.

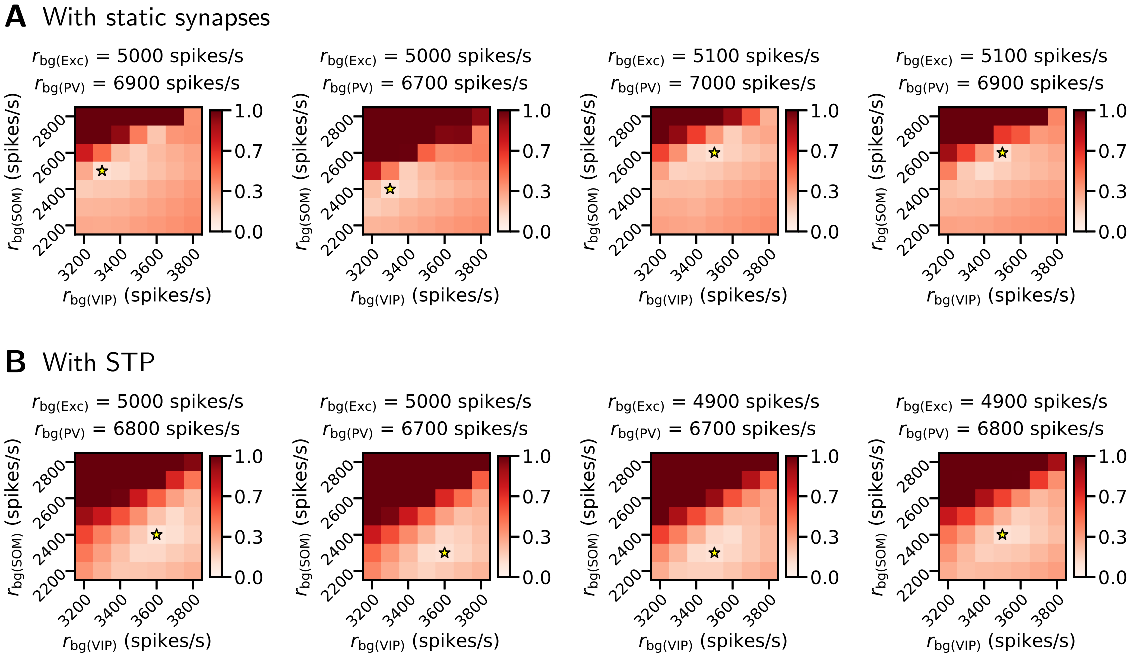


**Figure S3: Root-mean-square error of L2/3 and L4 relative population firing rates between simulation and experiment in models with different firing rates of cell-type-specific background inputs.** $r_{\mathrm{bg(Exc)}}$, $r_{\mathrm{bg(PV)}}$, $r_{\mathrm{bg(SOM)}}$, $r_{\mathrm{bg(VIP)}}$: firing rates of background inputs for Exc, PV, SOM, VIP cells, respectively. The star in each heatmap shows its best-fit point.

### Double-Sized Model

This model is adapted from the Base and Base-STP models by doubling the neuron number of each population and deriving the recurrent connection probabilities with twice the original area. Figure S4 shows the resting states, and Figures S5 and S6 show the results of cell-type-specific stimulation of this model. These results qualitatively reproduce those of the Base and Base-STP models with two exceptions, seen in Figure S5D. First, in the static synapse version of this model, L2/3 SOM cells undergo an initial increase in firing rate (albeit with large variability) upon VIP stimulation, whereas this stimulation always suppressed the L2/3 SOM cells in the Base model. Second, the Exc cells are first suppressed and then activated in response to VIP stimulation, while those in the Base model are always activated.

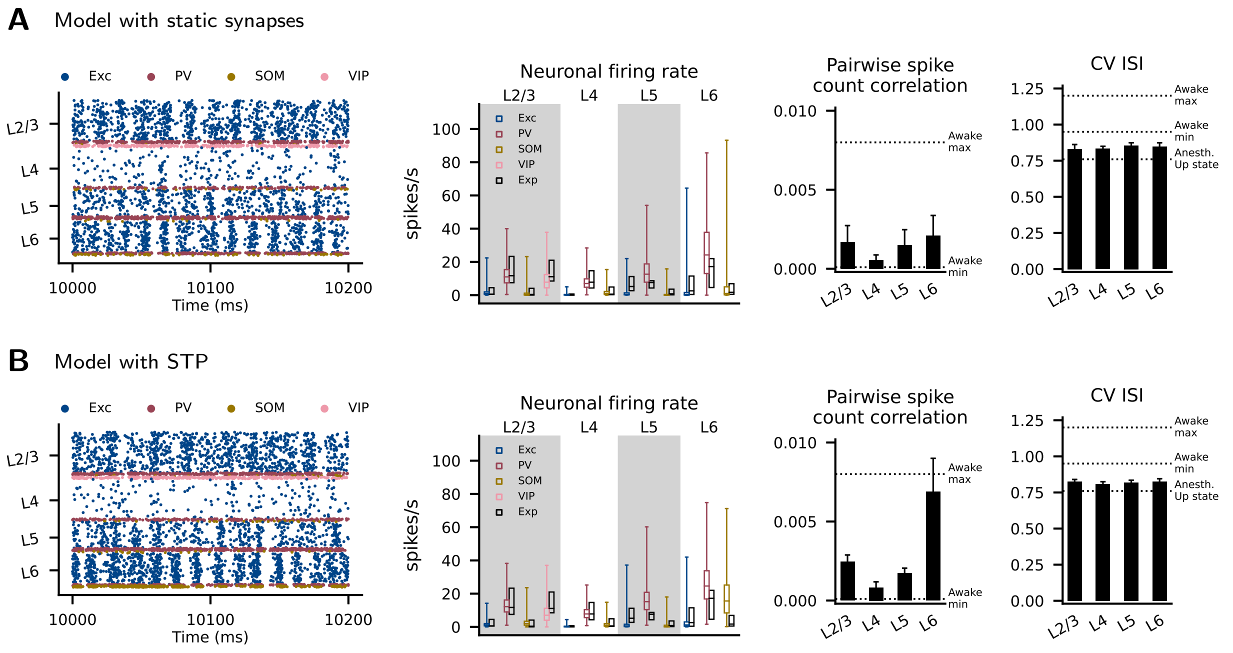


**Figure S4: Resting state of the double-sized model.** (A) The model version with static synapses. (B) The model version with STP. Left to right: raster plot of neuronal firings, neuronal firing rate of each cortical population, pairwise spike count correlation in each layer, and coefficients of variation of inter-spike intervals (CV ISI) in each layer. For the neuronal firing rate, the colored boxplots show the medians, 25th and 75th percentiles, and ranges of all neurons from 20 simulation instances. The black boxplots show the corresponding experimental data from Table 1 in Yu et al. (2019). For the pairwise spike count correlation and CV ISI, the black bars show means and standard deviations across 20 simulations, and the dashed lines show the *in vivo* criteria as follows. Awake max, Awake min: the maximum and minimum in the means of 13 recording sessions in frontal cortices of awake rats. Anesth. Up state: the mean of recorded Up states in motor cortex neurons of anesthetized rats. These values were provided by Maksimov et al. (2018).

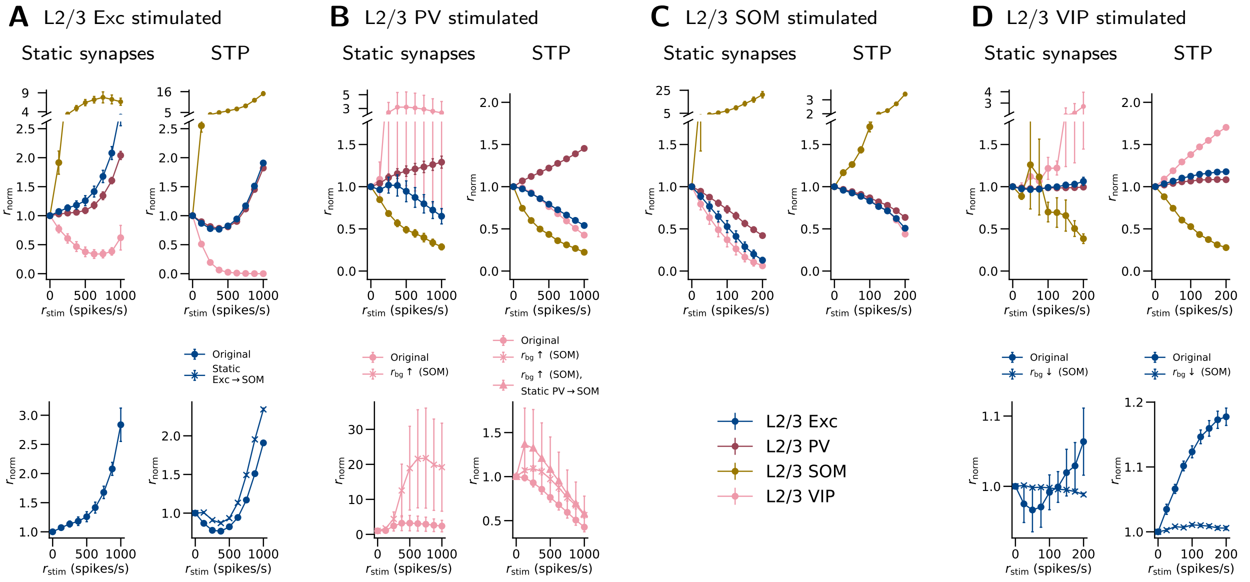


**Figure S5: L2/3 network responses of the double-sized model to cell-type-specific stimulation.** Each of (A) to (D) shows the changes in population firing rates when one particular cell type is stimulated in L2/3. The first row shows the results of the model where only the size is doubled as described ('Original'), while the second row compares with responses of further modified model versions, which reveals effects involving interneurons and their STP. $r_{\mathrm{stim}}$: rate of stimulation applied. $r_{\mathrm{norm}}$: resulting relative population firing rate, normalized to the data at $r_{\mathrm{stim}} = 0$. Original: original double-sized model. Static X→Y: STP in X→Y connections is excluded, X and Y being two cell types. $r_{\mathrm{bg}}\uparrow$ (X), $r_{\mathrm{bg}}\downarrow$ (X): $r_{\mathrm{bg}}$ for cell type X is increased or decreased. Error bars: SEM across simulation instances (n=20) with different randomization seeds. In some cases, the SEM is so small that the bar is not visible. The stimulation protocol is the same as in Figure 7 in the main article, except that the $r_{\mathrm{bg}}\uparrow$ (SOM) for STP in (B) represents that the $r_{\mathrm{bg}}$ for SOM is 100 spikes/s higher than in the 'Original' case, instead of 175 spikes/s higher.

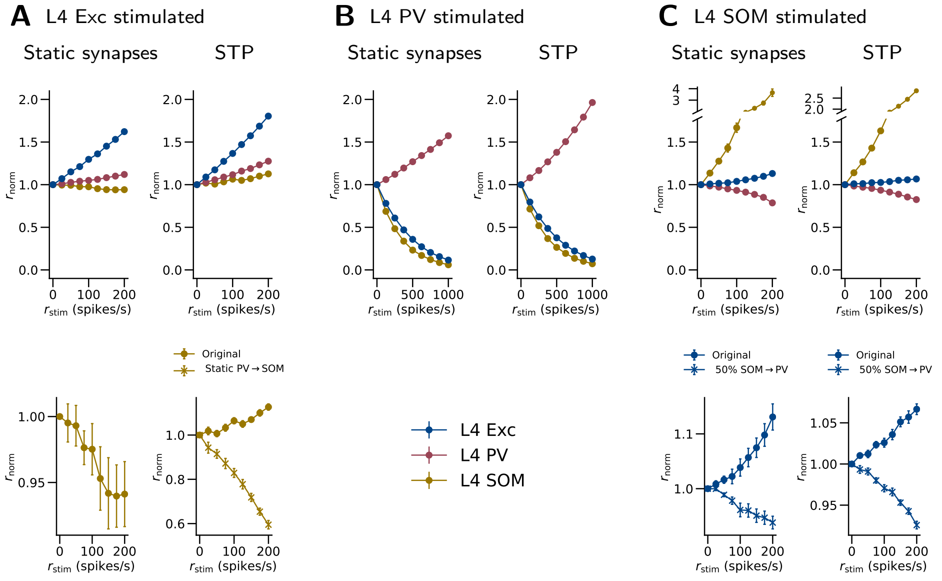


**Figure S6: L4 network responses of the double-sized model to cell-type-specific stimulation.** The notations are as in Figure S5. The stimulation protocol is the same as in Figure 8 in the main article.

### TC-Adjusted Model

This model is adapted from the Base and Base-STP models by downscaling the excitatory synaptic weights of recurrent connections to L2/3 Exc, SOM, and VIP cells (see Network Responses to Thalamic Stimulation). Figure S7 shows the resting states, and Figures S8 and S9 show the results of cell-type-specific stimulation of this model. These results qualitatively reproduce those of the Base and Base-STP models with two exceptions. In the STP version of this model, the Exc cells do not show a paradoxical decrease in firing rate when stimulated (Figure S8A), and excluding the STP in PV→SOM connections does not produce an initial VIP activation seen in the Base-STP model (Figure S8B).

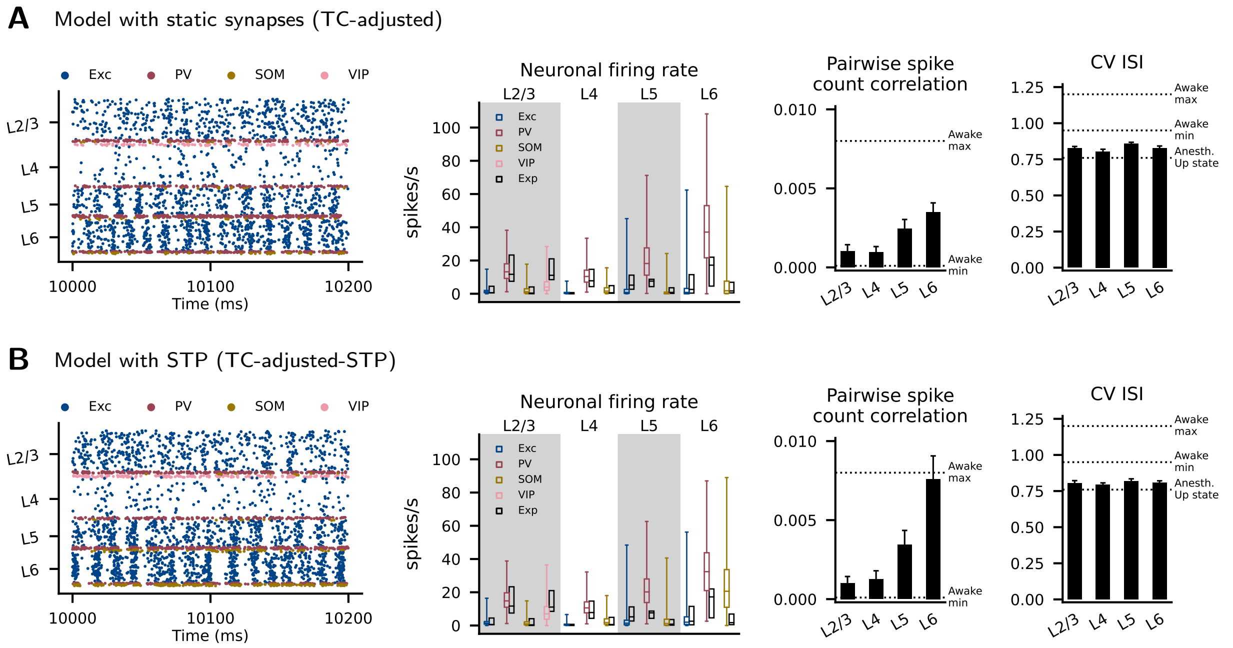


**Figure S7: Resting state of the model adjusted to obtain more realistic thalamocortical responses.** (A) The model version with static synapses (TC-adjusted). (A) The model version with STP (TC-adjusted-STP). The notations are as in Figure S4.

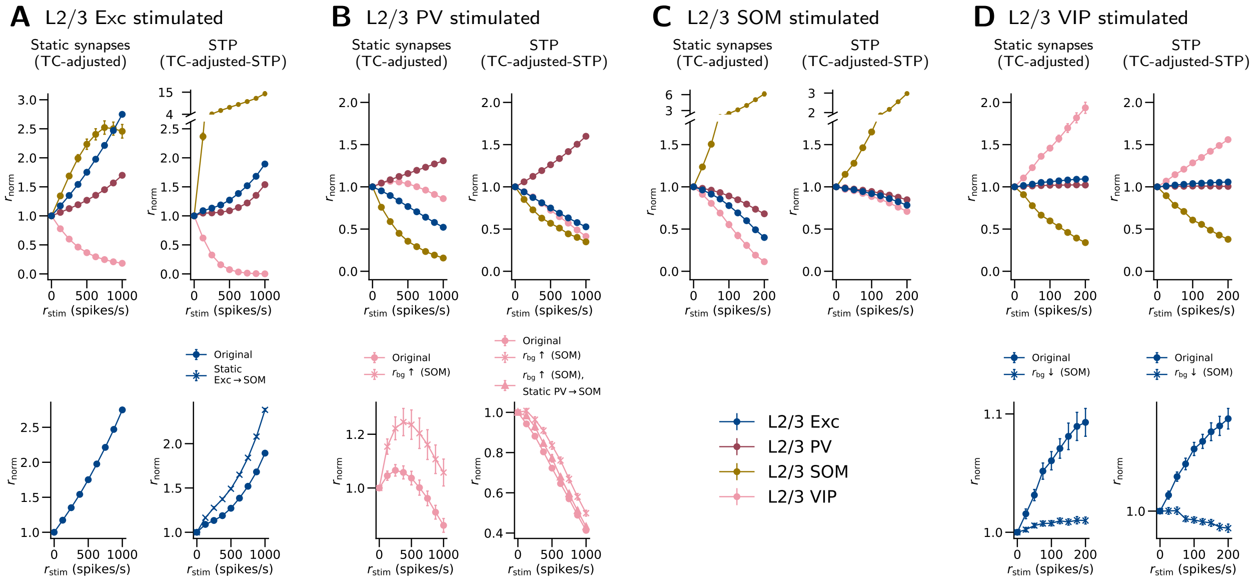


**Figure S8: L2/3 network responses of the TC-adjusted model and the TC-adjusted-STP model to cell-type-specific stimulation.** The notations are as in Figure S5. The stimulation protocol is the same as in Figure 7 in the main article.

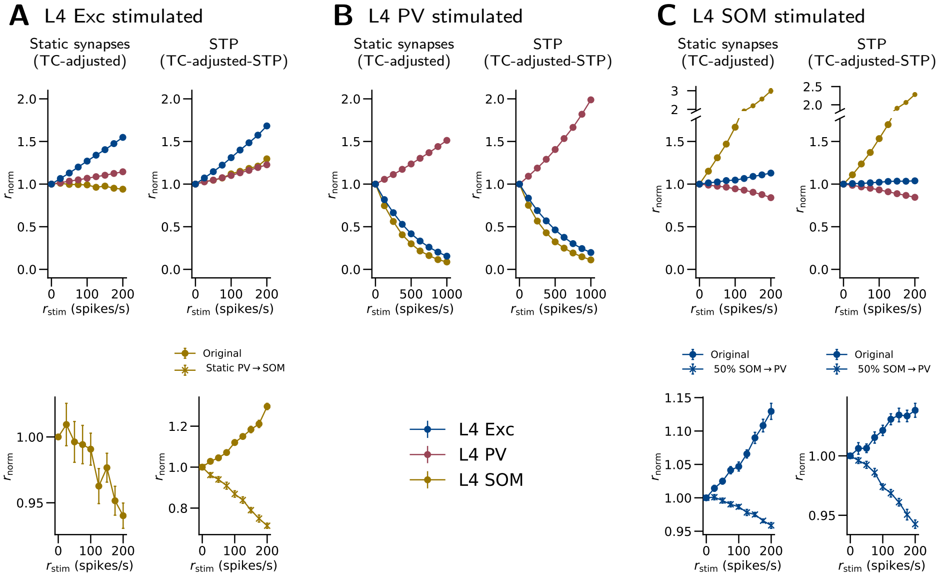


**Figure S9: L4 network responses of the TC-adjusted model and the TC-adjusted-STP model to cell-type-specific stimulation.** The notations are as in Figure S5. The stimulation protocol is the same as in Figure 8 in the main article.

### Second- and Third-Best-Fit Models in the Optimization of the Background Input

This section presents the second- and third-best-fit models in the parameter scans described in Optimization of the Background Input. These model versions are the same as the Base and Base-STP models except for the firing rates of the background inputs. Figures S10 and S11 show the resting states, and Figures S12, S13, S14, and S15 show the results of cell-type-specific stimulation of these model versions. These results qualitatively reproduce those of the Base and Base-STP models, except that the average L6 pairwise spike count correlation in the STP version of the second-best-fit model is slightly beyond the *in vivo* range (Figure S10B).

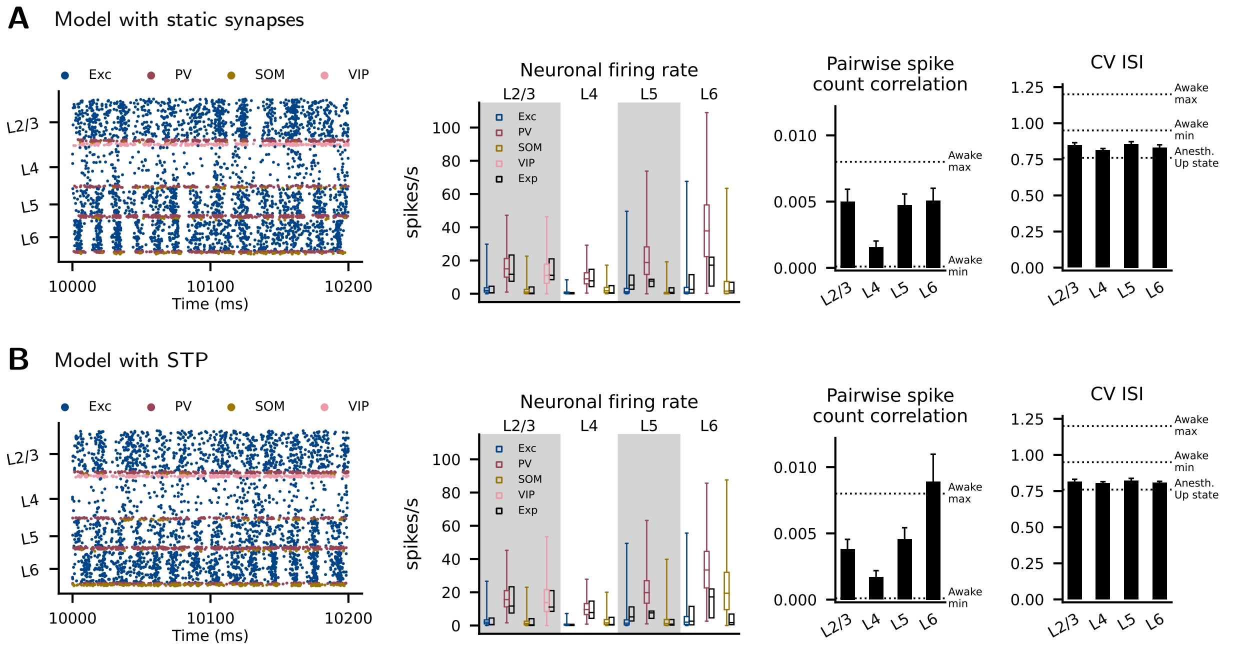


**Figure S10: Resting state of the second-best-fit model in the optimization of the background input.** (A) The model version with static synapses. (B) The model version with STP. The firing rates of background input for the model with static synapses are 5000 (Exc), 6700 (PV), 2400 (SOM), and 3300 (VIP) spikes/s. Those for the model with STP are 5000 (Exc), 6700 (PV), 2300 (SOM), and 3600 (VIP) spikes/s. The notations are as in Figure S4.

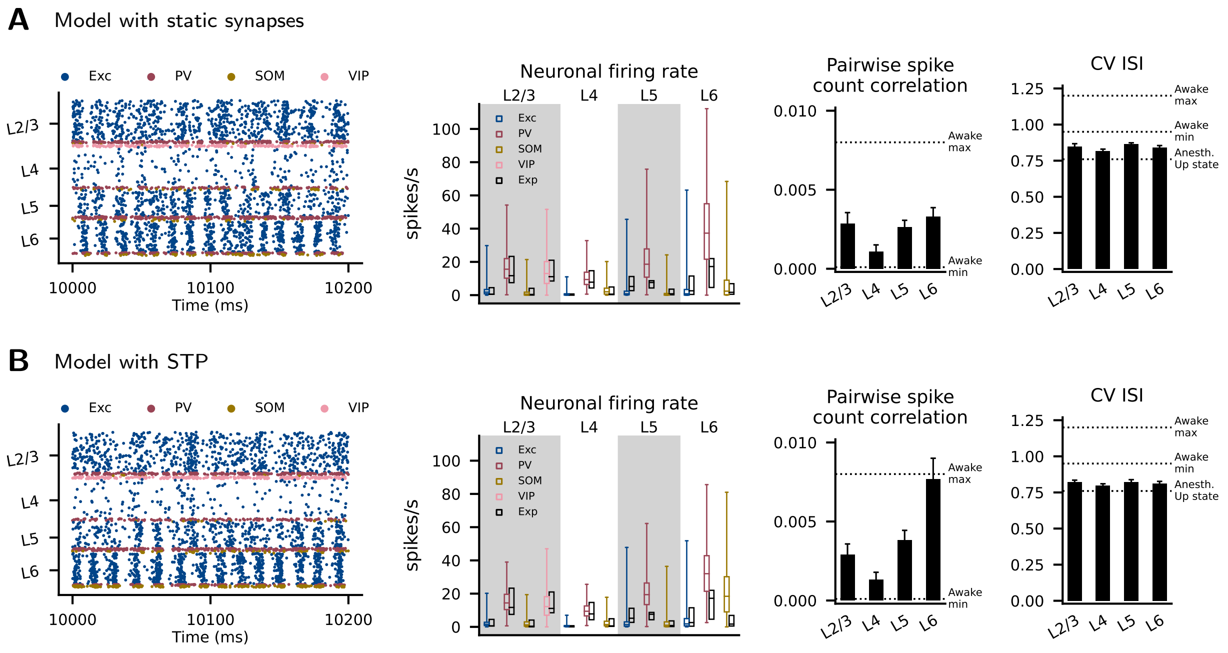


**Figure S11: Resting state of the model with the third-best-fit model in the optimization of the background input.** (A) The model version with static synapses. (B) The model version with STP. The firing rates of background input for the model with static synapses are 5100 (Exc), 7000 (PV), 2600 (SOM), and 3500 (VIP) spikes/s. Those for the model with STP are 4900 (Exc), 6700 (PV), 2300 (SOM), and 3500 (VIP) spikes/s. The notations are as in Figure S4.

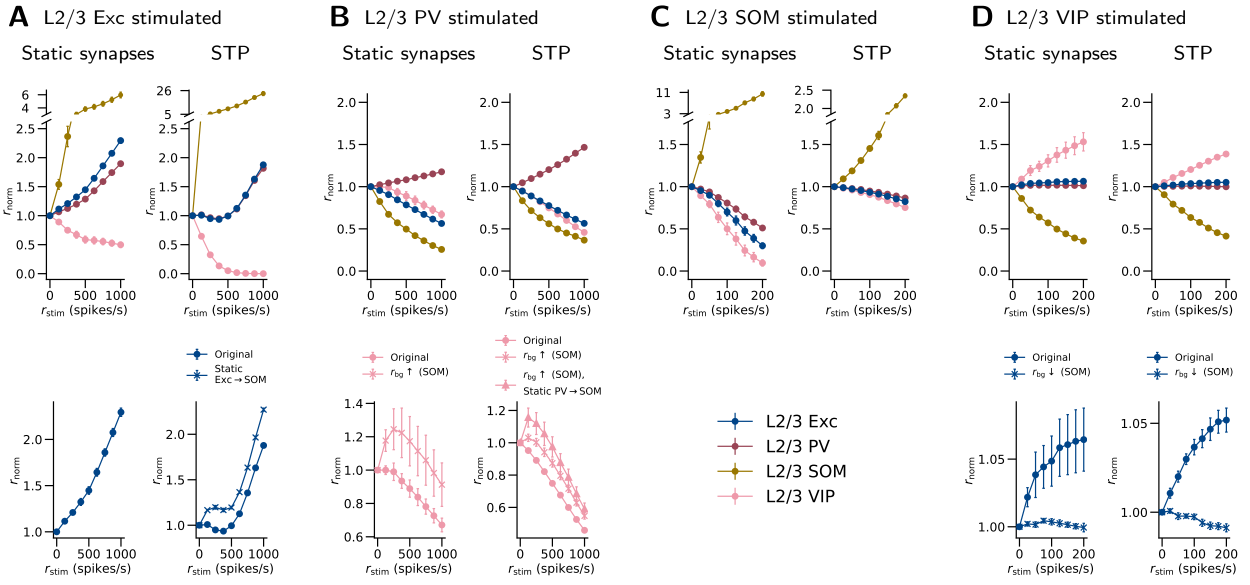


**Figure S12: L2/3 network responses to cell-type-specific stimulation of the second-best-fit model in the optimization of the background input.** The notations are as in Figure S5. The stimulation protocol is the same as in Figure 7 in the main article, except that the $r_{bg}\uparrow$ (SOM) for STP in (B) represents that the $r_{bg}$ for SOM is 200 spikes/s higher than in the 'Original' case, instead of 175 spikes/s higher.

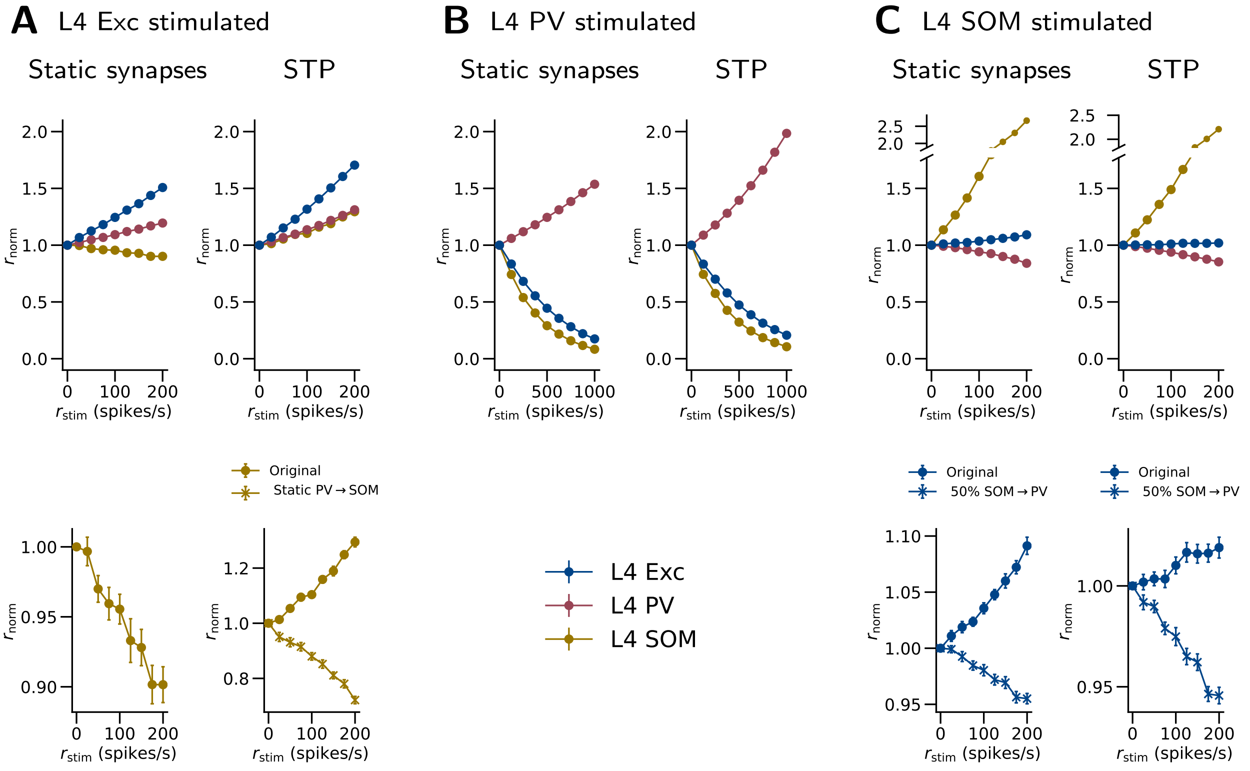


**Figure S13: L4 network responses to cell-type-specific stimulation of the second-best-fit model in the optimization of the background input.** The notations are as in Figure S5. The stimulation protocol is the same as in Figure 8.

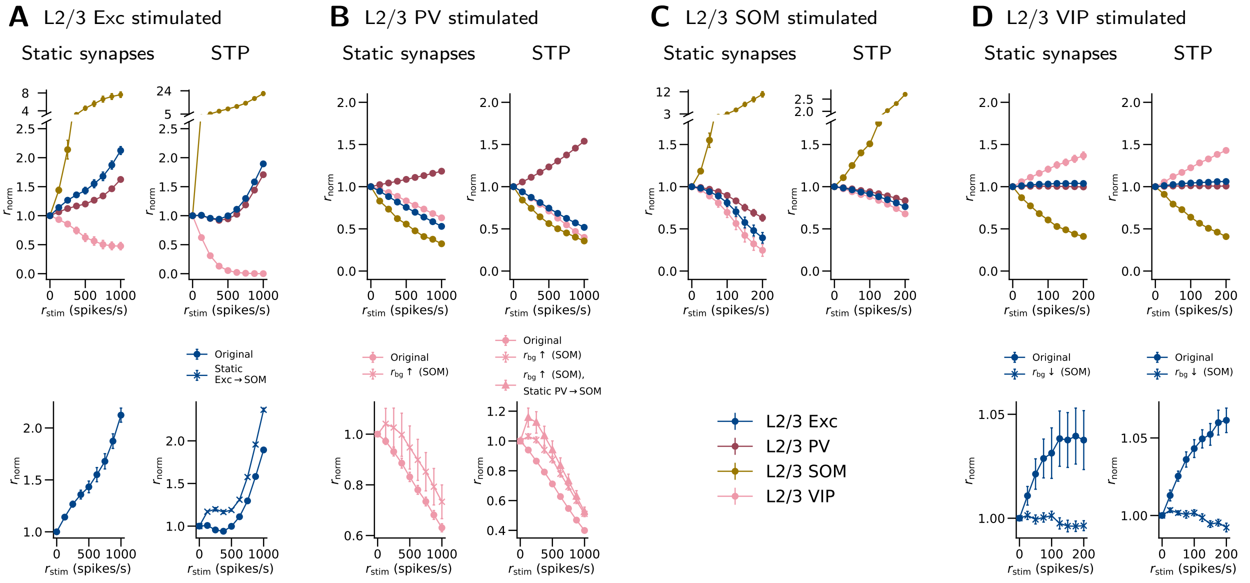


**Figure S14: L2/3 network responses to cell-type-specific stimulation of the third-best-fit model in the optimization of the background input.** The notations are as in Figure S5. The stimulation protocol is the same as in Figure 7 in the main article, except that the $r_{bg}$↑ (SOM) for STP in (B) represents that the $r_{bg}$ for SOM is 200 spikes/s higher than in the 'Original' case, instead of 175 spikes/s higher.

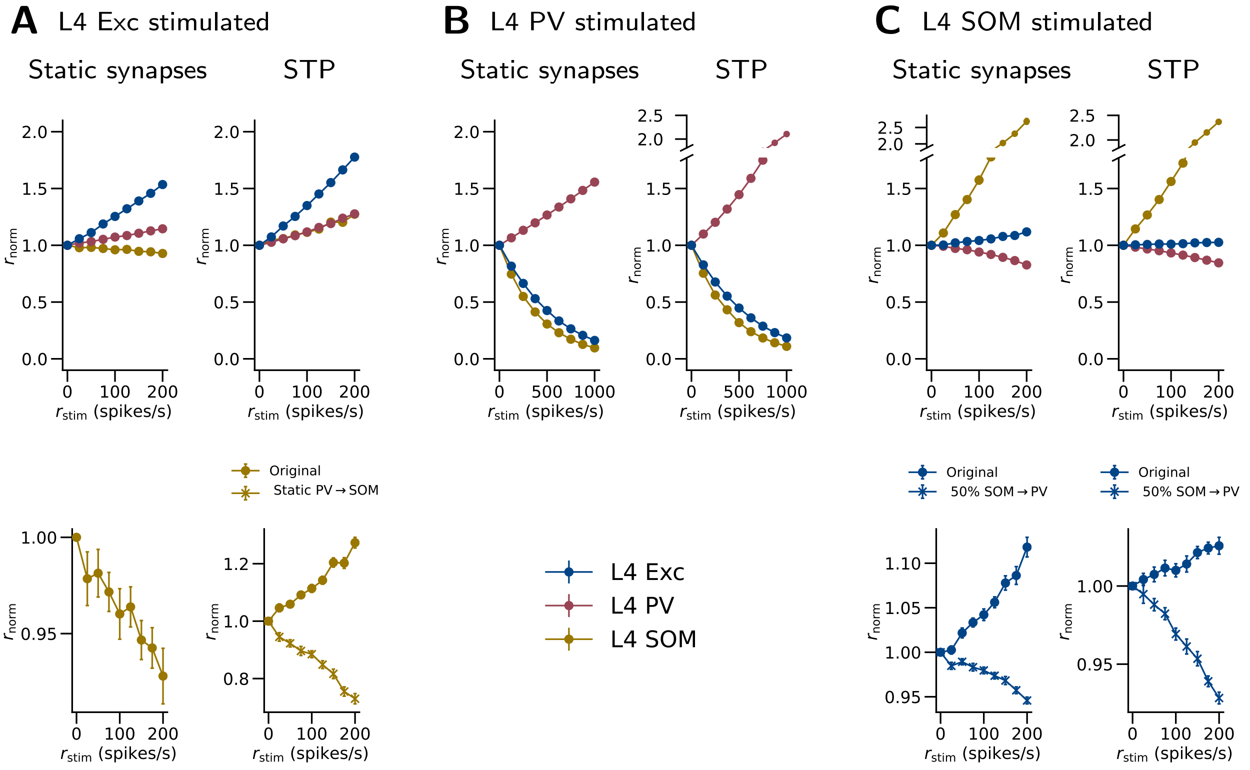


**Figure S15: L4 network responses to cell-type-specific stimulation of the third-best-fit model in the optimization of the background input.** The notations are as in Figure S5. The stimulation protocol is the same as in Figure 8 in the main article.

### Examination of the Paradoxical Effect in PV Cells

In this section, we examine the paradoxical effect in PV cells. This effect refers to the experimental observation that PV cells show an initial paradoxical decrease in activity when stimulated with excitatory inputs (Mahrach et al., 2020; Sanzeni et al., 2020). Here we examine if this paradoxical effect can be reproduced with several versions of our model (with static synapses). This is done with cell-type-specific stimulation of L2/3 PV cells as described in the main article, but with weaker stimulus strengths.

The paradoxical effect is not observed in PV cells in the Base model (Figure S16A) or in models of this size with modified parameters (Figure S16B to G). In contrast, the double-sized model (Figure S16H) shows a slight paradoxical decrease in L2/3 PV activity at low stimulation levels ($-1.2\%$ at $r_{\mathrm{stim}} = 12.5$ spikes/s and $-1.0\%$ at $r_{\mathrm{stim}} = 25$ spikes/s).

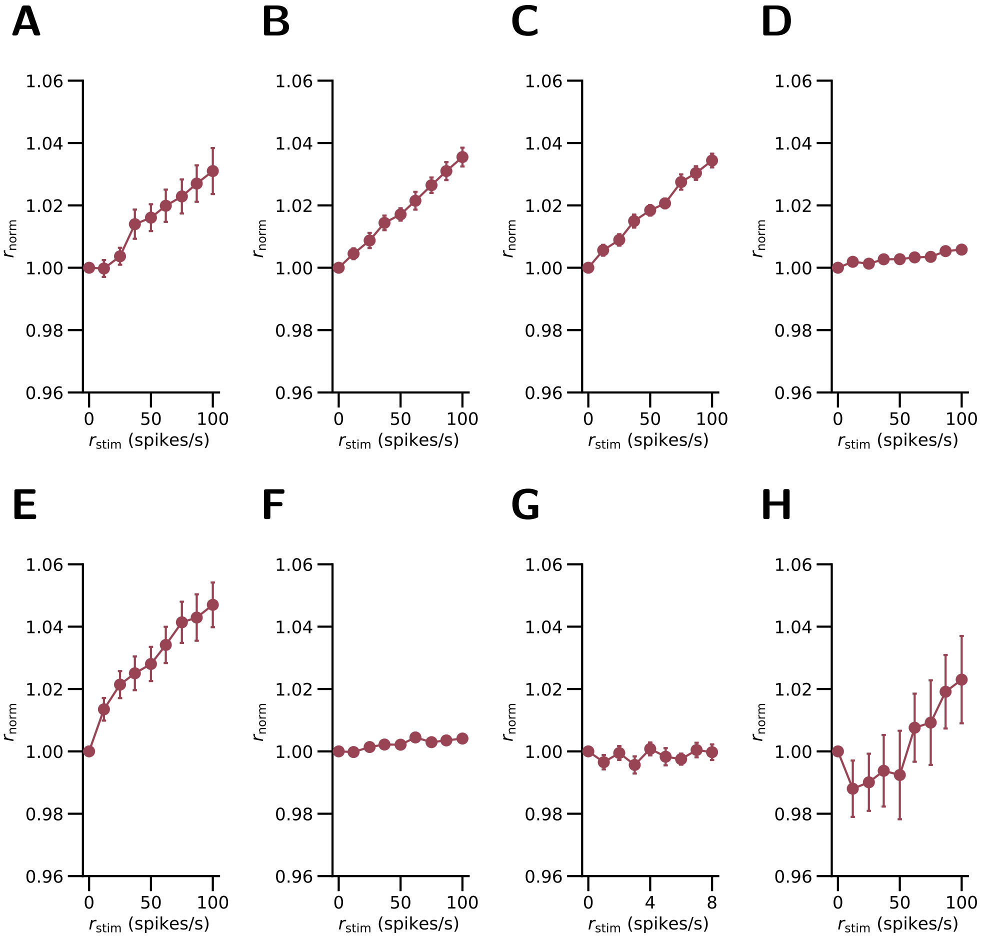


**Figure S16: Changes in L2/3 PV population firing rates in response to Poisson inputs.** (A) The Base model. (B) L2/3 Exc→Exc connections are removed. (C) Probability of the L2/3 Exc→Exc connections is 1/128 of that in the original model. (D) VIP→Exc, VIP→PV, and VIP→VIP connections in L2/3 are removed. (E) L4 to L6 of the model is removed. (F) Combining (D) and (E). (G) The stimulation rates ($r_{\mathrm{stim}}$) are lower (0 to 8 spikes/s). (H) The double-sized model. $r_{\mathrm{norm}}$: resulting relative population firing rate, normalized to the data at $r_{\mathrm{stim}} = 0$. The stimulation protocol is the same as in Figure 7 in the main article but with different levels of $r_{\mathrm{stim}}$.

### Importance of the Recurrent and Background Inputs

Figure S17 compares the resting-state population firing rates under conditions where a certain part of the connectivity in the model is removed. When the intralaminar connections are removed, the firing rates are much higher than in the original (Base) model. When the interlaminar connections are removed, the change is relatively small.

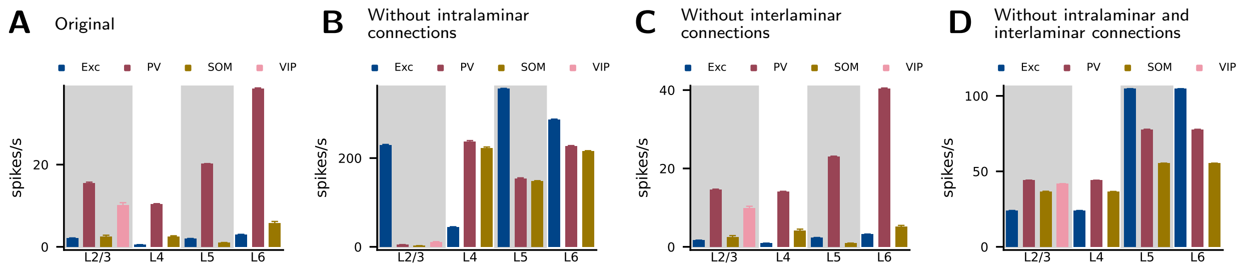


**Figure S17: Population firing rates in the original model and the model versions with modified connectivity.** (A) Original model (Base). (B) Without intralaminar connections. (C) Without interlaminar connections. (D) Without intralaminar and interlaminar connections. Error bars: standard deviations across 20 simulation instances.
